# Novel synthetic ecteinascidins exhibit potent anti-melanoma activity by suppressing super-enhancer-driven oncogenic transcription

**DOI:** 10.1101/2024.03.26.586754

**Authors:** Max Cigrang, Julian Obid, Maguelone Nogaret, Léane Seno, Tao Ye, Guillaume Davidson, Philippe Catez, Pietro Berico, Clara Capelli, Clara Marechal, Amélie Zachayus, Clémence Elly, Tsai-Kun Li, Emmanuel Compe, Pablo Avilés, Irwin Davidson, Jean-Marc Egly, Carmen Cuevas, Frédéric Coin

**Author notes:** Co-first authors (Max Cigrang, Julian Obid).

## Abstract

The dynamic cellular transitions exhibited by skin cutaneous melanoma (SKCM) cells present a significant challenge to current therapeutic approaches, emphasizing the critical need for innovative treatments. Lurbinectedin, a marine-derived compound belonging to the ecteinascidin family, has recently gained approval for the treatment of metastatic small-cell lung cancer (SCLC). In this study, we demonstrate the efficacy of lurbinectedin against SKCM cells, irrespective of their driver mutations or phenotypic states. Additionally, we have developed two novel derivatives of lurbinectedin, termed ecubectedin and PM54, both of which exhibit potent cytotoxic effects on SKCM cells. Moreover, these analogs demonstrate robust anti-tumor activity in melanoma xenograft models, including those resistant to current therapies, leading to prolonged animal survival. Mechanistically, our investigation reveals that these novel synthetic ecteinascidins markedly suppress oncogenic super-enhancer (SE)-mediated gene expression in SKCM cells through a multifaceted mechanism. They bind to and inhibit the activity of promoters of lineage-specific master transcription factors, as well as promoters of genes encoding ubiquitous transcription factors/coactivators, which are highly enriched at oncogenic SEs. These mechanisms likely synergize to disrupt the expression of cancer-promoting genes. Overall, our findings highlight the potential of synthetic ecteinascidins as promising therapeutics for cancers characterized by diverse transcriptional landscapes, particularly in cases where conventional therapeutic options have failed due to the heterogeneity of malignant cell population

## Introduction

Malignant skin cutaneous melanoma (SKCM), comprising only 1 % of skin cancer cases, is responsible for 80 % of related deaths (NCI-SEER-Database, 2023) ^1, 2^. SKCM has been pointed to as a prime example of how the understanding of biological mechanisms can be translated into novel therapeutics ^3, 4, 5^. Comparative genomic studies have identified key targetable driver mutations in SKCM, with aberrant activation of the Mitogen Activated Protein Kinase (MAPK) pathway observed in 90 % of cases due to somatic mutations in the BRAF (50 %), RAS (20 %) and NF1 (15 %) oncogenes ^6, 7^. Patients with the commonly found BRAF^V600E/K^ mutations, leading to constitutive MEK and ERK signaling, can benefit from combined treatment with targeted BRAF or MEK therapies, resulting in favorable progression-free survival rates ^8, 9, 10, 11, 12, 13, 14^. Additionally, immune checkpoint inhibitors targeting CTLA-4 and PD-1 have become the first-line treatment for metastatic melanoma, providing long-term benefits to a significant number of patients ^15, 16^.

Despite the advancements in targeted and immune-therapies, complete remission is achieved only in a small subset of patients, while severe adverse effects and limited efficacy are observed in a majority of cases ^10, 17, 18, 19^. Moreover, non-BRAF mutated melanoma poses significant challenges, as effective treatment options are limited ^20^. One of the critical barriers to clinical success is intrinsic or acquired insensitivity to treatment. Various mechanisms of drug resistance have been described, with intra-tumoral heterogeneity driven by cellular phenotypic plasticity emerging as a key contributor to relapse ^21, 22, 23, 24, 25^. Indeed, melanoma cells can undergo phenotype switching, transitioning between melanocytic/differentiated states governed by genes with essential roles in cell proliferation such as the lineage-specific master transcription factor MITF, and mesenchymal-like/undifferentiated states governed by the key regulators AXL and AP-1/TEAD genes, implicated in drug resistance and invasion ^26, 27, 28, 29, 30, 31, 32, 33^. As such, phenotypic adaptation, rendered possible through dynamic transcriptional and epigenetic reprogramming mechanisms in response to microenvironmental cues, complicates treatment outcomes ^34, 35^. The heterogeneity and phenotypic plasticity of melanoma cells underscore the need for therapeutics that can uniformly target divergent transcription programs governing different tumor cell states ^36^.

In recent years, the concept of ‘transcriptional addiction’ has gained attention as a novel hallmark of cancer cells. Dysregulated gene expression programs and their associated transcriptional regulatory machinery are critical for sustaining cancer cell phenotypes, making them susceptible to transcriptional inhibitors ^37, 38, 39, 40, 41^. One of the main mechanisms leading to gene expression dysregulation in cancer cells consists in the abnormal acquisition of large clusters of enhancers known as “super-enhancers” (SEs), driving and maintaining the robust expression of oncogenes. SEs are characterized by higher levels of the Mediator coactivator complex and aggregated histone modifications H3K27ac and H3K4me1 over longer genomic distances, compared to typical enhancers ^42, 43, 37^. Furthermore, SE-dependent oncogene transcription requires the activity of ubiquitous transcription factors (such as the CDK7 kinase of TFIIH) and transcriptional coactivators (such as Bromodomain-containing protein 4 (BRD4)). Disruption of these oncogenic SEs emerges as a potential therapeutic option. Therefore, several compounds targeting factors involved in oncogenic SE-driven gene expression, including CDK7 and BRD4 inhibitors, have entered clinical trials ^44, 45^, however with limited successes so far due to poor pharmacokinetics and short half-lives ^46, 47, 48, 49^.

Lurbinectedin, a synthetic analog of marine-derived ecteinascidins, is a DNA binder recently approved for the treatment of relapsed small cell lung cancer (SCLC) in several countries, including the USA and Canada ^50, 51, 52^. In an effort to potentially further enhance the benefits of lurbinectedin, we developed ecubectedin and PM54, two new analogs derived from lurbinectedin. Our study initially focused on analyzing the sensitivity to lurbinectedin, ecubectedin and PM54 of a diverse panel of human SKCM cell lines and cell cultures, which are characterized by different oncogenic alterations and cellular states, and therefore different degrees of resistance to currently used anti-melanoma therapies. We first demonstrated potent anti-proliferative and apoptotic effects of these new molecules on differentiated and undifferentiated BRAF, NRAS, and triple-wild type mutated melanoma cells in various *in vitro* 2-D and 3-D models and *in vivo* in cell-derived xenograft (CDX) mouse models. Secondly, we sought to better understand the precise mechanism of action of these novel types of anti-cancer molecules. Through chemical-mapping approaches, direct observation of transcriptional condensates in melanoma cell nuclei and transcriptomic studies in CDX tumors, we further discovered that the new synthetic ecteinascidins potently disrupt oncogenic SE-driven transcription through a multifaceted mechanism. We observed that these compounds distinguish themselves from traditional chemotherapies by exclusively binding to transcriptionally active, preferentially CG-rich genomic regions. As such, strong drug-binding was observed in promoter regions of genes encoding ubiquitous transcription factors/coactivators such as CDK7, CDK12, EP300 or BRD4, heavily enriched at SEs, thereby inhibiting their expression and consequently decommissioning oncogenic SEs. Genes encoding lineage-specific master transcription factors such as MITF or FOSL2, known to be involved in SE-maintaining auto-regulatory circuits, were also found to be strongly bound and inhibited. Synthetic ecteinascidins likely disrupt the formation of phase-separated condensates at SEs themselves, by directly binding to these regulatory elements. Interestingly, our studies suggest potential advantages of PM54 over the parent compound, with a restricted set of genes inhibited by this drug compared to lurbinectedin, while retaining equivalent efficacy. This multifaceted mode of action ensures potent disruption of oncogenic transcription. Intriguingly, our data suggest that these newly uncovered mechanisms are also at work in SCLC cells, upon treatment with synthetic ecteinascidins. These findings provide a compelling rationale for investigating ecteinascidins in clinical trials for treatment-resistant melanoma and pave the way for further research and development of these compounds as effective treatment for other cancers characterized by diverse transcriptional landscapes and in which conventional therapies have failed.

## Results

### Lurbinectedin exhibits notable efficacy against distinct melanoma cell types

To investigate the response of melanoma cells to lurbinectedin (**Figure 1a**), we examined ten different melanoma cells representing the two primary phenotypes and encompassing the most prevalent driver mutations in SKCM. On one hand, we assessed differentiated patient-derived melanocytic-type cultures, including MM011 (NRAS^Q61K^), MM074 (BRAF^V600E^), MM117 (Triple-wt), alongside melanoma cell lines 501mel (BRAF^V600E^), IGR37 (BRAF^V600E^) and SkMel-28 (BRAF^V600E^). These cells exhibited moderate to high expression levels of the lineage-specific master transcription factors MITF and SOX10, while demonstrating low to undetectable expression levels of the pro-metastatic factors EGFR and AXL ^53, 30, 47^ (**Figure 1b and Table 1**). Conversely, we examined patient-derived undifferentiated and mesenchymal-like melanoma cell cultures MM029 (BRAF^V600K^), MM047 (NRAS^Q61R^), MM099 (BRAF^V600E^) and the melanoma cell line IGR39 (BRAF^V600E^). These cells displayed low to undetectable levels of MITF and SOX10, but elevated expression levels of EGFR and/or AXL ^47^ (**Figure 1b and Table 1**).

**Figure 1:**
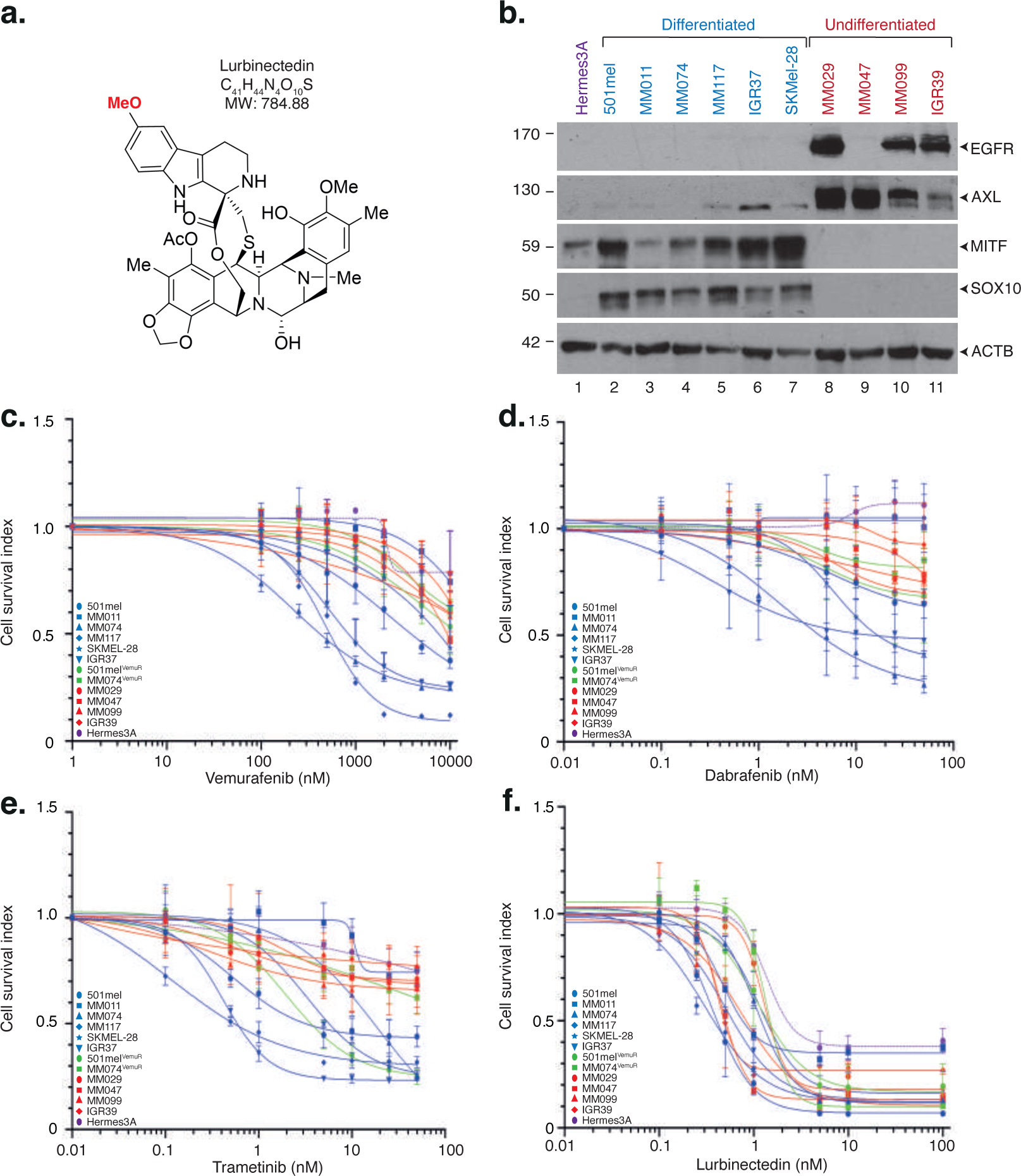
Melanoma cells show high sensitivity to lurbinectedin. **a.** Chemical structure of lurbinectedin, a synthetic ecteinascidin containing tetrahydroisoquinoline subunits. The moiety of the molecule interacting with DNA binding is indicated. Molecular Weight (MW) is indicated. **b.** Protein lysates from either the immortalized Hermes3A melanocytes, differentiated melanoma cells 501mel, MM011, MM074, MM117, IGR37 and SKMel-28 or undifferentiated melanoma cells MM029, MM047, MM099 and IGR39 were immuno-blotted for proteins as indicated. Molecular mass of the proteins is indicated (kDa). **c-f.** Melanoma cells were treated with increasing concentrations of vemurafenib **(c)**, dabrafenib **(d)**, trametinib **(e)**, lurbinectedin **(f)** for 72 hours. Mean growth is shown relative to vehicle (DMSO)-treated cells. Error bars indicate mean values +/− Standard Deviation (SD) for three biological triplicates. Differentiated (MITF-High, proliferative) melanoma cells are shown in blue, while undifferentiated (MITF-low, invasive) melanoma cells are shown in red. Differentiated melanoma cells with acquired resistance to Vemu are shown in green. Immortalized Hermes3A melanocytes are shown in violet.

**Table 1:** Melanoma cell sensitivity against MAPKi and synthetic ecteinascidins. IC50 of vemurafenib, dabrafenib, trametinib, lurbinectedin, ecubectedin and PM54 against various melanoma cells. The phenotype and genotype of these cells are indicated. Hermes3A are transformed melanocytes.

Using cell viability assay, we observed that the patient-derived cell cultures and melanoma cell lines exhibited varying sensitivities to targeted therapy agents such as the BRAF inhibitors (BRAFi) Vemurafenib and Dabrafenib, as well as the MEK inhibitor (MEKi) Trametinib (**Figures 1c-e and Table 1**). Differentiated BRAF^V600E^ melanoma cells, such as MM074 or IGR37, were the most responsive to these compounds, while undifferentiated cells demonstrated high resistance. In contrast, we observed that all melanoma cells displayed high sensitivity to Lurbinectedin, with IC50 (half maximal inhibitory concentration) values in the low nanomolar range, spanning from 0.44 to 2.07nM (**Figure 1f and Table 1**). Additionally, we generated Vemurafenib-resistant cells, namely 501mel^VemuR^ and MM074^VemuR^, by exposing initially sensitive cells to increasing drug concentrations *in vitro* (**Figure 1c and Table 1**) ^47^. These Vemurafenib-resistant cells acquired a hyperpigmentation phenotype ^47^ and exhibited cross-resistance to Dabrafenib (in the case of MM074^VemuR^) and Trametinib (**Figures 1d-e and Table 1**), but remained highly sensitive to Lurbinectedin (**Figure 1f and Table 1**). Strikingly, we observed that the non-cancerous Hermes3A immortalized melanocytes were consistently 3- to 7-times less sensitive than the melanoma cells towards Lurbinectedin.

Collectively, these findings underscore the heightened sensitivity of melanoma cells to Lurbinectedin, irrespective of the cellular phenotypes or driver mutations.

### Two novel ecteinascidins show high cytotoxic effect on melanoma cells

In our pursuit of enhancing the anti-cancer efficacy of novel compounds, we synthesized and assessed two derivative molecules closely related to lurbinectedin. These compounds, named ecubectedin and PM54, exhibit distinct chemical structures in the moieties of the molecules that are not engaged in DNA binding ^54^: Ecubectedin features a substituted spiro β-carboline, while PM54 contains a spiro benzofuropyridine-a moiety not previously identified in ecteinascidins (**Figures 2a-b**). These structural variations may confer unique pharmacological properties, warranting further investigation. Using cell viability assay, we observed that all melanoma cells displayed high sensitivity to these new ecteinascidins, with IC50 values falling within the low nanomolar range, spanning from 0.7 to 5nM (**Figures 2c-d and Table 1**). In addition to proliferative and undifferentiated states, single-cell sequencing has in recent years unveiled the existence of additional cell states within melanoma tumors, such as interferon-active melanoma cells ^35, 33^. The significance of these cells in the context of resistance to treatment has been significantly underestimated. To generate pseudo-interferon-active melanoma cells, we subjected MM074 cells to treatment with interferon-γ, resulting in the expression of *bona fide* markers of the interferon-active state such as PD-L1, IRF1 or STAT1 and its active phosphorylated form, pSTAT1 (**Supplemental Figure 1a**). Notably, this induction was accompanied by the acquisition of resistance to BRAFi (**Supplemental Figure 1b and Table 1**). Intriguingly, these pseudo-interferon-active melanoma cells exhibited sustained sensitivity to synthetic ecteinascidins (**Supplemental Figure 1c and Table 1**).

**Figure 2:**
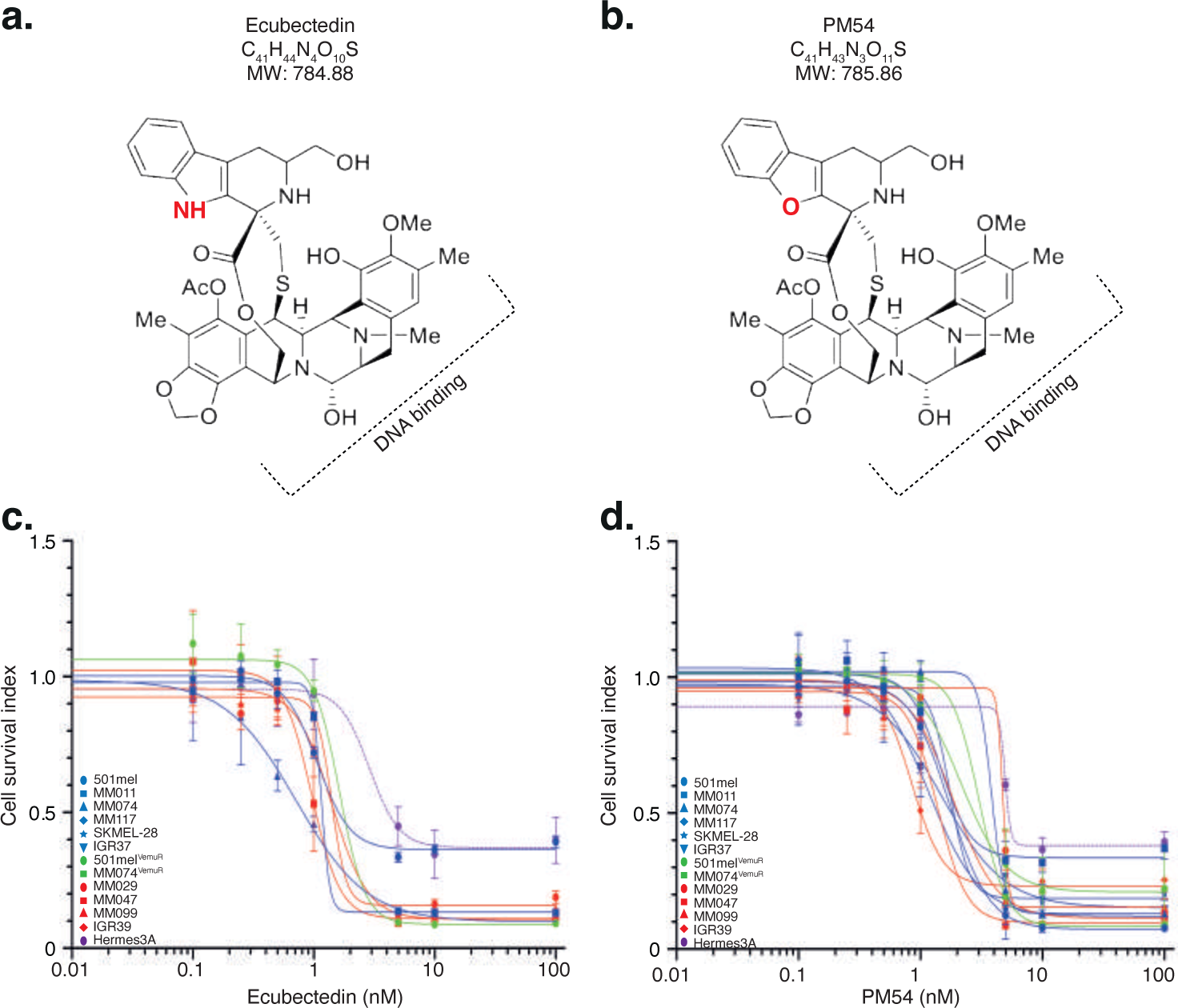
Melanoma cells show high sensitivity to novel synthetic ecteinascidins. **a-b.** Chemical structure of the novel ecteinascidin analogs ecubectedin **(a)** and PM54 **(b)**, derived from lurbinectedin. The modifications are highlighted in red. The moiety of the molecule interacting with DNA binding is indicated. Molecular Weight (MW) is indicated. **c-d.** Melanoma cells were treated with increasing concentrations of ecubectedin **(c)** or PM54 **(d)**, for 72 hours. Mean growth is shown relative to vehicle (DMSO)-treated cells. Error bars indicate mean values +/− Standard Deviation (SD) for three biological triplicates. Differentiated (MITF-High, proliferative) melanoma cells are shown in blue, while undifferentiated (MITF-low, invasive) melanoma cells are shown in red. Differentiated melanoma cells with acquired resistance to vemurafenib are shown in green. Immortalized Hermes3A melanocytes are shown in violet.

These findings clearly demonstrate that the two newly synthetized ecteinascidin analogs exhibit cytotoxic effects comparable to lurbinectedin on a range of melanoma cells containing distinct driver mutations and cellular phenotypes.

### Synthetic ecteinascidins induce melanoma cell apoptotic death

We next compared the efficacy of synthetic ecteinascidins on melanoma cell proliferation and survival. Initially, clonogenic assays demonstrated a significant impact of these molecules on all tested melanoma cell cultures or cell lines (**Figure 3a**) together with a significant inhibition of melanoma cell proliferation (**Figure 3b**). Concurrently, there was a notable blockade of cell cycle progression (**Figure 3c**) and induction of apoptosis (**Figure 3d**). We also observed that synthetic ecteinascidins significantly affected the invasion (**Figure 3e**) and migration (**Figure 3f**) of undifferentiated melanoma cell cultures.

**Figure 3:**
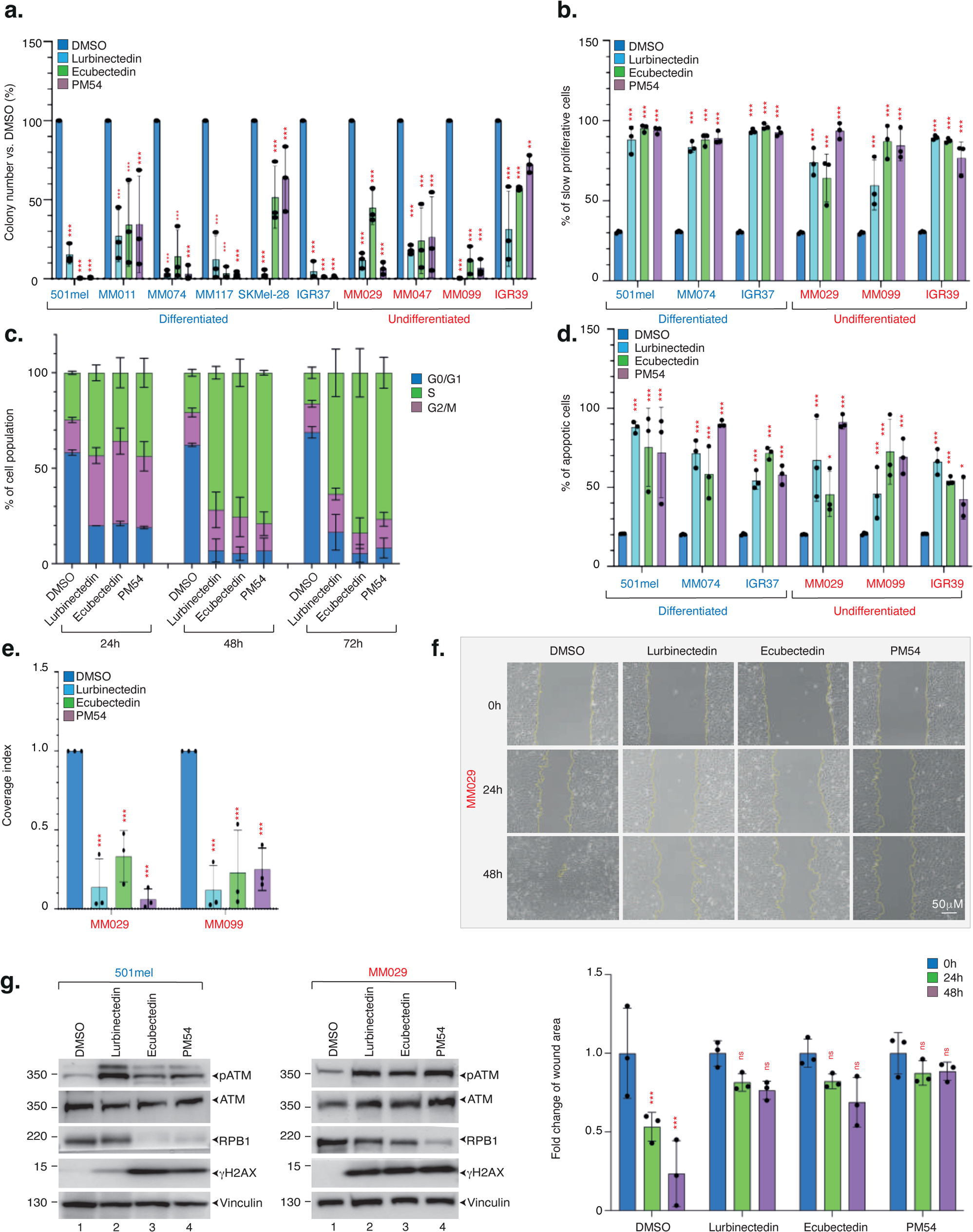
Synthetic ecteinascidins induce cell cycle arrest and apoptosis. **a.** Indicated melanoma cells were treated with either vehicle (DMSO), lurbinectedin, ecubectedin or PM54 (1xIC50 concentration, 48 hours) and then allowed to grow for additional 10 days in the absence of the drugs. Results are shown as the mean colony numbers +/− SD for three biological triplicates. Ordinary one-way ANOVA using Dunnett’s multiple comparisons test was used to determine the p-values (*vs.* DMSO). **b.** Indicated melanoma cells were incubated with CellTrace and subsequently treated with either vehicle (DMSO), lurbinectedin, ecubectedin or PM54 (1xIC50 concentration, 72 hours). Quantifications of populations with high CellTrace signal in DMSO or drug-treated cells are shown as mean values +/− SD for three biological triplicates. Proliferative cells show low CellTrace signal while non proliferative cells show high CellTrace signal. Ordinary one-way ANOVA using Dunnett’s multiple comparisons test was used to determine the p-values (*vs.* DMSO). **c.** 501mel cells were treated with either vehicle (DMSO), lurbinectedin, ecubectedin or PM54 (1xIC50 concentration, 72 hours). Cell cycle was studied by propidium iodide staining and flow cytometry, and results are shown as mean values +/− SD for three biological triplicates. **d.** Indicated melanoma cells were treated with either vehicle (DMSO), lurbinectedin, ecubectedin or PM54 (1xIC50 concentration, 72 hours). Apoptosis was studied by flow cytometry using annexin V-APC staining. Results are shown as mean values +/− SD for three biological triplicates. Ordinary one-way ANOVA using Dunnett’s multiple comparisons test was used to determine the p-values (*vs.* DMSO). **e.** MM029 and MM099 melanoma cells were treated with either vehicle (DMSO), lurbinectedin, ecubectedin or PM54 (1xIC50 concentration,48 hours). Invasion was determined using Boyden chamber assays. Results are shown as mean values of coverage index +/− SD for three biological triplicates. Ordinary one-way ANOVA using Dunnett’s multiple comparisons test was used to determine the p-values (*vs.* DMSO). **f.** Confluent monolayers of MM029 cells were scratched and fresh medium containing reduced FCS % and either vehicle (DMSO), lurbinectedin, ecubectedin or PM54 (1xIC50 concentration) was added (left). Size of the wound was measured at the indicated times and results are shown as mean values of fold changes of wound area *vs.* DMSO treatment +/− SD for three biological triplicates (right). Ordinary one-way ANOVA using Dunnett’s multiple comparisons test was used to determine the p-values (*vs.* DMSO). **g.** Protein lysates from differentiated 501mel or undifferentiated MM029 melanoma cell, as indicated, treated with either lurbinectedin, ecubectedin or PM54 (5xIC50 concentration, 24 hours) were immuno-blotted for proteins as indicated. Molecular mass of the proteins is indicated (kDa).

In SCLC models, lurbinectedin induces the degradation of the largest subunit of RNA Polymerase II (RPB1) and triggers a DNA damage response characterized by the activation of γH2AX due to drug-induced DNA breaks ^50, 51^. Using immunofluorescence, we observed γH2AX accumulation in the nuclei of differentiated 501mel melanoma cells or undifferentiated MM029 cell cultures upon treatment with synthetic ecteinascidins (**Supplemental Figures 2a-b-c-d**), which was confirmed by immunoblotting in differentiated 501mel and MM074 cells or undifferentiated MM029 and MM099 cells (**Figure 3g and Supplemental Figure 2e**). In parallel, phosphorylation of ATM, the master damage response protein, was observed in differentiated 501mel and undifferentiated MM029 cells (**Figure 3g**). Remarkably, while lurbinectedin induced minimal RPB1 degradation, the presence of ecubectedin and PM54 resulted in highly pronounced degradation of RPB1 in these cells, highlighting the superior efficacy of the new compounds in this context (**Figure 3g**).

We next employed melanosphere culture assays to investigate the impact of synthetic ecteinascidins on three-dimension (3-D) melanoma cultures. Initially, we assessed the response of melanospheres derived from the melanocytic-like MM074 cells to BRAFi and MEKi. In sharp contrast to the response observed in 2-D cultures, BRAFi and MEKi failed to reduce cell viability in 3-D culture, even at doses equivalent to 5x of the IC50 determined in 2-D (**Supplemental Figure 3a**). Conversely, synthetic ecteinascidins demonstrated significant cytotoxic effects on MM074 melanospheres at nanomolar concentrations (**Supplemental Figures 3b-c**).

These findings elucidate the potent cytostatic and cytotoxic impacts of synthetic ecteinascidins on both differentiated and undifferentiated melanoma cells, marked by the induction of DNA breaks and the degradation of RNAPII. It is noteworthy that this impact is especially prominent in the case of ecubectedin and PM54.

### Synthetic ecteinascidins exhibit robust anti-tumor activities

The above data prompted us to examine the impact of synthetic ecteinascidins *in vivo* on melanoma cell-derived xenograft (CDX) mouse models. We first monitored the tumor volumes following intravenous (IV) administration of synthetic ecteinascidins once per week for three consecutive weeks at a concentration of 1.2mg/kg. Treatments commenced (d.0) when the tumors reached 150 mm^3^ in athymic nude female aged 4 to 6 weeks (N=8/group), and finished fourteen days later (d.14). We tested CDXs obtained from two highly proliferative melanoma cell lines widely used for drug screening (LOX-IMVI^BRAF-V600E^ and WM-266-4^BRAF-V600D^) ^55^. For both CDXs, we observed significant tumor growth regression upon treatment with synthetic ecteinascidins, starting d.5. The tumor growth delay was persistent even after d.14 when treatment was withdrawn, and last until d.25, emphasizing a period of latency of 10 days following the end of the treatment. Simultaneously, a marked augmentation in overall survival was observed, predominantly evident during the latency phase (**Figures 4a-c**).

**Figure 4:**
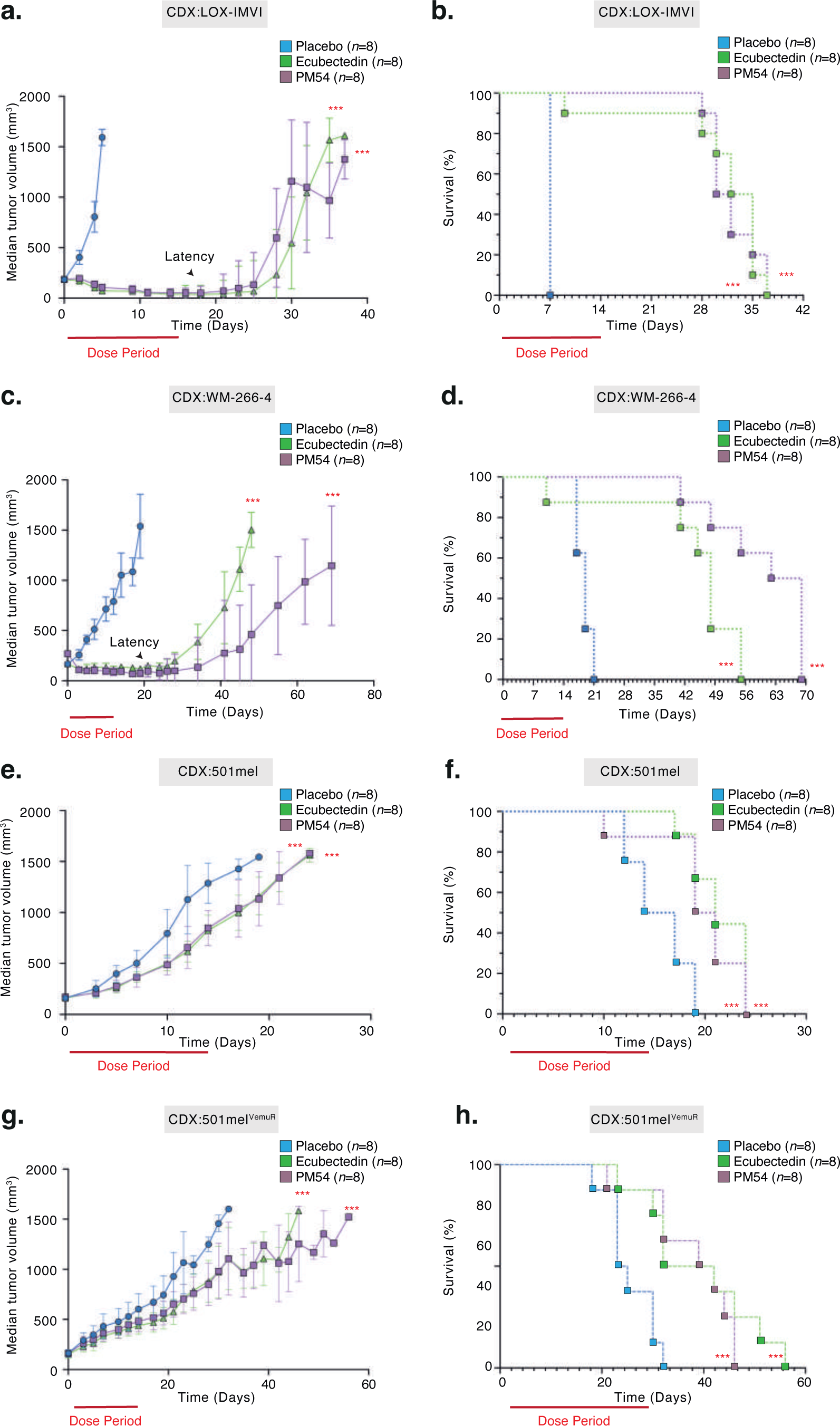
Potent *in vivo* effects of synthetic ecteinascidins. **a-c-e-g.** Indicated CDX models (n=8) were treated with Placebo, ecubectedin or PM54 at 1.2 mg/kg once a week for 3 consecutive weeks (on days 0, 7 and 14) and tumor volumes were measured. Red bar indicates the dose period. The latency phase is indicated by an arrow. Logrank (Mantel-Cox) test was used to determine the p-values. **b-d-f-h**. Indicated CDX models (n=8) were treated weekly with Placebo, ecubectedin or PM54 at 1.2 mg/kg and survival was assessed. Red bar indicates the dose period. The latency phase is indicated by an arrow. Logrank (Mantel-Cox) test was used to determine the p-values.

We next analyzed the effect of the drugs on MAPKi-resistant cells using the 501mel and 501mel^VemuR^ cells **(see Figure 1**). Once the tumors reached a size of 150 mm^3^ in female NSG mice, a single IV dose of either ecubectedin or PM54 at a concentration of 1.2mg/kg was administrated to the animals (N=8/group). Twenty-four hours after this single IV dose, we assessed both the mitotic and apoptotic indexes using immunostaining of phospho-histone H3 (pHH3) and caspase-3 cleavage, respectively ^56^, on tumor sections. We observed a significantly decreased mitotic index and increased apoptosis upon treatment with synthetic ecteinascidins, for both CDXs derived from 501mel and 501mel^VemuR^ (**Supplemental Figures 4a-d**). Consequently, we observed that treatments with synthetic ecteinascidins impacted the tumor growth of CDXs derived from 501mel and 501mel^VemuR^ melanoma cells (**Figure 4 e-h**).

Altogether, these studies suggest that synthetic ecteinascidins are highly active at inhibiting the growth of melanoma tumors, even those presenting resistance to clinically relevant treatments.

### PM54 differentially affects expression of genes in melanoma cells

Given the established impact of lurbinectedin on transcription in SCLC ^51^, we conducted gene expression profiling (RNA-seq) within 2-D cultures of differentiated and undifferentiated melanoma cells (MM074 and MM029, respectively). Following the treatment with lurbinectedin, a significant down-regulation of 2,357 in differentiated cells and 2,757 genes in undifferentiated cells was observed (**Figure 5a and Supplemental Table 1)**. Another subset of genes, specifically 1,219 in differentiated cells and 1,968 in undifferentiated cells, exhibited up-regulation in response to treatment. For ecubectedin and PM54, a significant down-regulation of 2,185 and 1,889 in differentiated cells and 2,820 and 2,083 genes in undifferentiated cells was observed, respectively **(Supplemental Table 1 and Figure 5b-c**). Again, a significantly fewer number of genes (more particularly for PM54) was up-regulated in these cells following ecubectedin or PM54 treatment (1,196 and 936 in differentiated cells and 2,046 and 409 genes in undifferentiated cells for ecubectedin and PM54, respectively). Among the down-regulated genes, we noted the presence of several lineage-specific master transcription factors and regulators such as MITF, PAX3 or SOX10 in differentiated melanoma cells or AXL, EGFR, SOX9, FOSL2 and TEAD4 in undifferentiated cells. These data were confirmed in 2-D models by RT-qPCR and/or immunoblotting (**Supplemental Figures 5a-d**) and in 3-D models by RT-qPCR (**Supplemental Figures 5e-f**).

**Figure 5:**
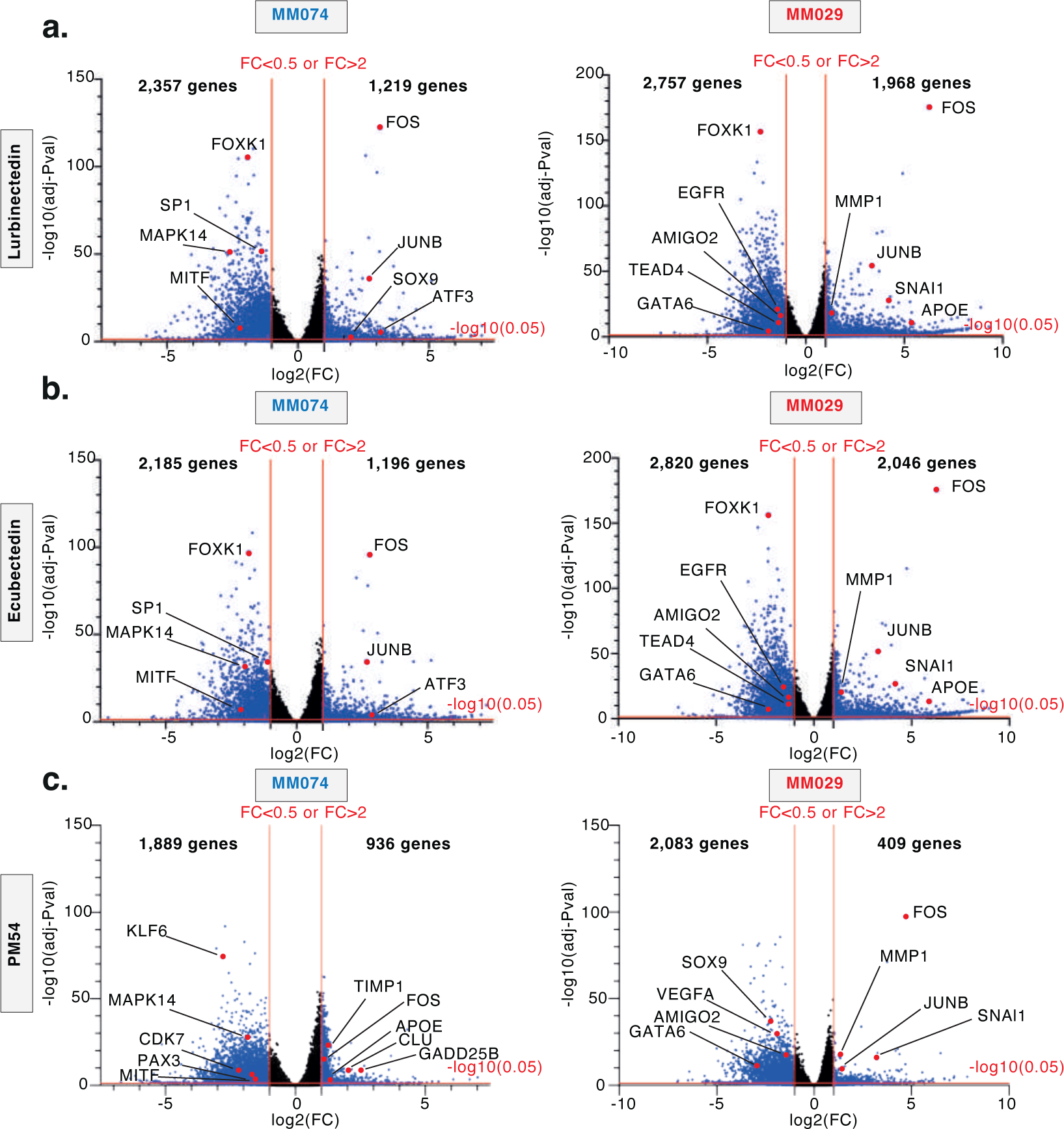
Synthetic ecteinascidins affect the transcription program of melanoma cells. **a-c**. Volcano plots showing differentially expressed genes between DMSO-treated *vs.* **(a)** Lurbi-, **(b)** Ecubectedin-, or **(c)** PM54-treated MM074 (left) or MM029 (right) cells, determined by RNA-seq (10xIC50 concentration, 8 hours). Examples of significantly deregulated genes are shown, which were defined as genes with log2(Fold change) > 1 or < −1 and adjusted P-value < 0.05.

We next undertook a comparative analysis of gene expression profiles in response to treatment with synthetic ecteinascidins. Notably, the three molecules commonly down-regulated 1,365 and 1,104 genes in differentiated and undifferentiated cells, respectively (**Supplemental Figure 6a**). It is worth mentioning that among these genes, 757 displayed consistent down-regulation (and only 110 displayed up-regulation) across both differentiated and undifferentiated cells in response to all the three compounds (**Supplemental Figure 6b and Supplemental Table 2)**. Gene ontology (GO) analysis revealed that a substantial proportion of these 757 genes were intricately involved in transcriptional processes (**Supplemental Figure 6c**).

We subsequently compared each novel synthetic ecteinascidin with lurbinectedin. We observed that ecubectedin exhibited strikingly similar effects, with no genes exhibiting statistically significant differential expression upon a comparative analysis in either differentiated or undifferentiated cells (**Supplemental Figure 7a and b**). In stark contrast, PM54 distinctly induced specific transcriptional effects compared to lurbinectedin, revealing a more focused alteration in gene expression, as a smaller subset of genes exhibited deregulation in both differentiated and undifferentiated melanoma cells, (**Supplemental Figure 7a and c**). This distinction was further substantiated by Gene Set Enrichment Analysis (GSEA) and GO analysis, which elucidated that PM54 exerts weaker effects on genes involved in diverse cellular processes such as interferon response or oxidative phosphorylation but exerts a more direct influence on genes involved in transcriptional regulation (**Supplemental Figures 8a and b**).

Collectively, our findings underscore the profound impact of synthetic ecteinascidins on the transcriptional programs within melanoma cells. Notably, PM54 distinguishes itself by exhibiting cytotoxic activity comparable to that of lurbinectedin and ecubectedin, yet remarkably, it exerts the least influence on the transcriptional program of melanoma cells, emphasizing a unique and potentially advantageous pharmacological profile.

### Coactivator condensation at SEs is sensitive to synthetic ecteinascidins

We next conducted a comprehensive analysis of RNA-seq datasets obtained after lurbinectedin treatment of cells from three different types of cancer (SKCM, SCLC and Non-SCLC). This analysis revealed that a common set of 642 genes underwent significant down-regulation upon drug exposure (**Figure 6a and Supplemental Table 3)**. GO analysis revealed a strong enrichment of genes involved in transcriptional regulation (**Supplemental Figure 9a**), with notable downregulated genes including ubiquitous transcription factors/coactivators (such as CDK7, CDK12, CDK13, EP300, CBP, BRD4) and Mediator complex subunits (such as CDK8 and MED13). These results were confirmed in differentiated and undifferentiated melanoma cells by immunoblotting (**Figure 6b and Supplemental Figure 9b**). Notably, *in vivo* experiments utilizing melanoma CDXs also demonstrated a rapid down-regulation of these genes, together with lineage-specific master transcription factors such as MITF, SOX10 or PAX3 following short-term treatment with ecteinascidins (**Figure 6c and Supplemental Figure 9c**).

**Figure 6:**
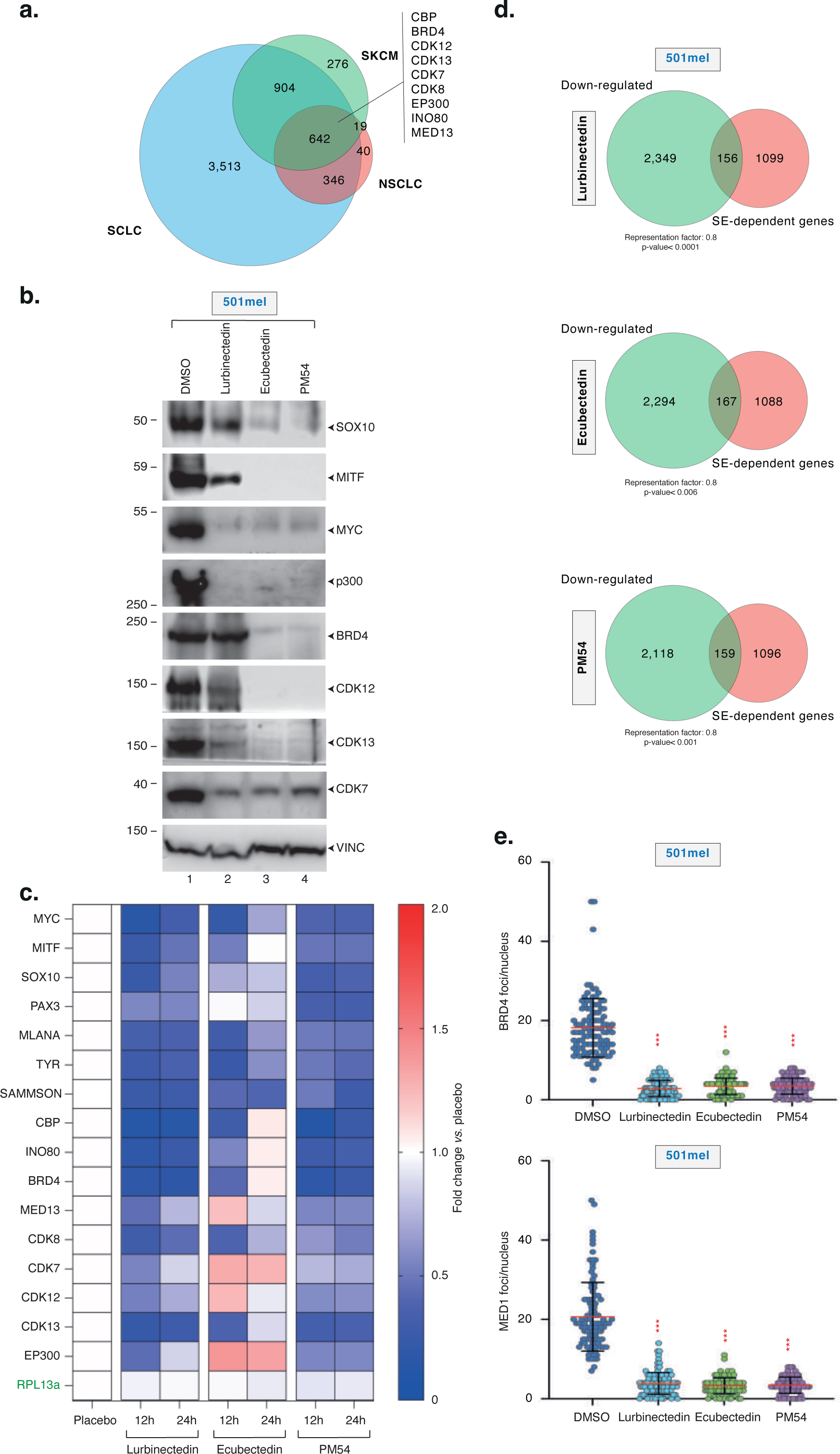
Synthetic ecteinascidins decommission SEs in melanoma cells. **a.** Venn diagram showing the overlap of genes down-regulated in SKCM (GSE256100), SCLC (GSE179074) and NSCLC (GSE179074), following treatment with Lurbinectedin. **b.** Differentiated 501mel melanoma cells were treated with ecteinascidins as indicated (5xIC50 concentration, 24 hours) and protein lysates were immuno-blotted for proteins as indicated. Molecular mass of the proteins is indicated (kDa). **c.** CDXs from 501mel cells (n=3) were treated with a single dose of lurbinectedin, ecubectedin or PM54 at 1.2 mg/kg and tumors were collected 12 or 24 hours later. Heatmap shows average placebo-normalized expression of the indicated genes obtained by qRT-PCR analysis. RPL13a is a housekeeping gene. **d.** Venn diagram showing the overlap of genes downregulated by ecteinascidins, as indicated, in 501mel cells (10xIC50 concentration, 8 hours) and SE-dependent genes identified in 501mel cells using H3K27ac-and BRD4-profiling by Cut&Tag and the ROSE algorythm ^76^. Representation factor and hypergeometric p-value are indicated. **e.** The numbers of BRD4 (top) and MED1 (bottom) foci per nucleus observed in 501mel cells following treatment with DMSO or ecteinascidins (5xIC50 concentration, 24 hours) are shown +/− SD. Red bars indicate mean integrated density. One-way ANOVA with post-hoc Tukey adjustment comparisons were used to determine the p-values (*vs.* DMSO).

Ubiquitous transcription factors/coactivators and the mediator complex are pivotal in driving oncogenic expression in cancer cells by activating, among others, genes dependent on SEs. Therefore, in an effort to identify SEs in our melanoma cell models, we performed Cut&Tag assays targeting H3K27ac and BRD4 in differentiated and undifferentiated cells (501mel and MM029, respectively). Using the Rank Ordering of Super-Enhancers (ROSE) algorithm and cross-referencing the list of SEs identified from the Cut&Tag on H3K27ac and that on BRD4, we identified 533 and 347 *bona fide* SEs in differentiated and undifferentiated cells, respectively (**Supplemental Figures 10a and Supplemental Table 4)**. Subsequently, we identified by ROSE 1,255 and 951 genes putatively regulated by these *bona fide* SEs, in differentiated and undifferentiated cells, respectively (**Supplemental Figures 10b and Supplemental Table 4)**. Although 261 SE-dependent genes were shared between differentiated and undifferentiated cells, most SE-dependent genes seemed to be cell-state-specific. We next crossed these data with the list of downregulated genes in both differentiated and undifferentiated melanoma cells following treatments with synthetic ecteinascidins and observed a significant enrichment of SE-dependent genes among those down-regulated genes (**Figure 6d and Supplemental Figure 10c**). This was also observed *in vivo*, where SE-dependent oncogenes such as SAMMSON or MYC were strongly downregulated (**Figure 6c**).

It was demonstrated that transcriptional coactivators such as BRD4 or the MED1 subunit of the Mediator complex may be visualized as discrete puncta in the nuclei of cells and that SEs associate with these puncta ^57^. Immunofluoresence revealed nuclear puncta for both BRD4 and MED1 in differentiated or undifferentiated melanoma cells (**Figure 6e and Supplemental Figures 11a-b**). Short-term treatments with synthetic ecteinascidins caused a reduction in the number of BRD4 and MED1 puncta in both types of cells, suggesting that transcriptional condensates formed at SEs are sensitive to treatment with synthetic ecteinascidins. ChIP/RT-qPCR further revealed that the levels of BRD4 and H3K27ac were strongly reduced at SEs regulating MITF or SOX10 in differentiated cells, or AXL or EGFR in undifferentiated cells, upon short-term treatment with synthetic ecteinascidins (**Supplemental Figures 11c-d**).

These results collectively support the notion that SEs regulating the expression of critical oncogenes are decommissioned by synthetic ecteinascidins.

### Synthetic ecteinascidins specifically target transcriptionally active, CG-rich genomic regions

We next sought to comprehensively map the genome-wide binding sites of synthetic ecteinascidins in melanoma cells. Using bioactive biotinylated versions of lurbinectedin and PM54 (Bio-lurbi and Bio-PM54), we conducted chemical-mapping ^58^ with three biological replicates per compound, using both differentiated or undifferentiated melanoma cells (501mel and MM029, respectively). Our analysis revealed approximately 30,000 drug-binding sites in differentiated and 15,000 in undifferentiated cells **(Supplemental Table 5)**, demonstrating high reproducibility with Spearman correlations exceeding 0.7 across triplicates (**Supplemental Figure 12a**).

Notably, approximately 75% of the identified drug-binding sites were found to be located in gene regions, with promoter (∼25-34%) and intronic (∼32-35%) binding frequencies being consistent for both Bio-lurbi and Bio-PM54, in both cell types (**Figure 7a and Supplemental Figure 12b**). Genome-wide, peaks of synthetic ecteinascidins predominantly co-localized with the transcriptionally active H3K27ac chromatin mark, RNAPII, BRD4 and positive ATAC-seq signals, and not with the repressive H3K27me3 chromatin mark (**Figure 7b and Supplemental Figure 12c**). Overall, we observed a highly significant correlation between drug-bound genes and genes down-regulated by the drugs (**Figure 7c and Supplemental Figure 12d**).

**Figure 7:**
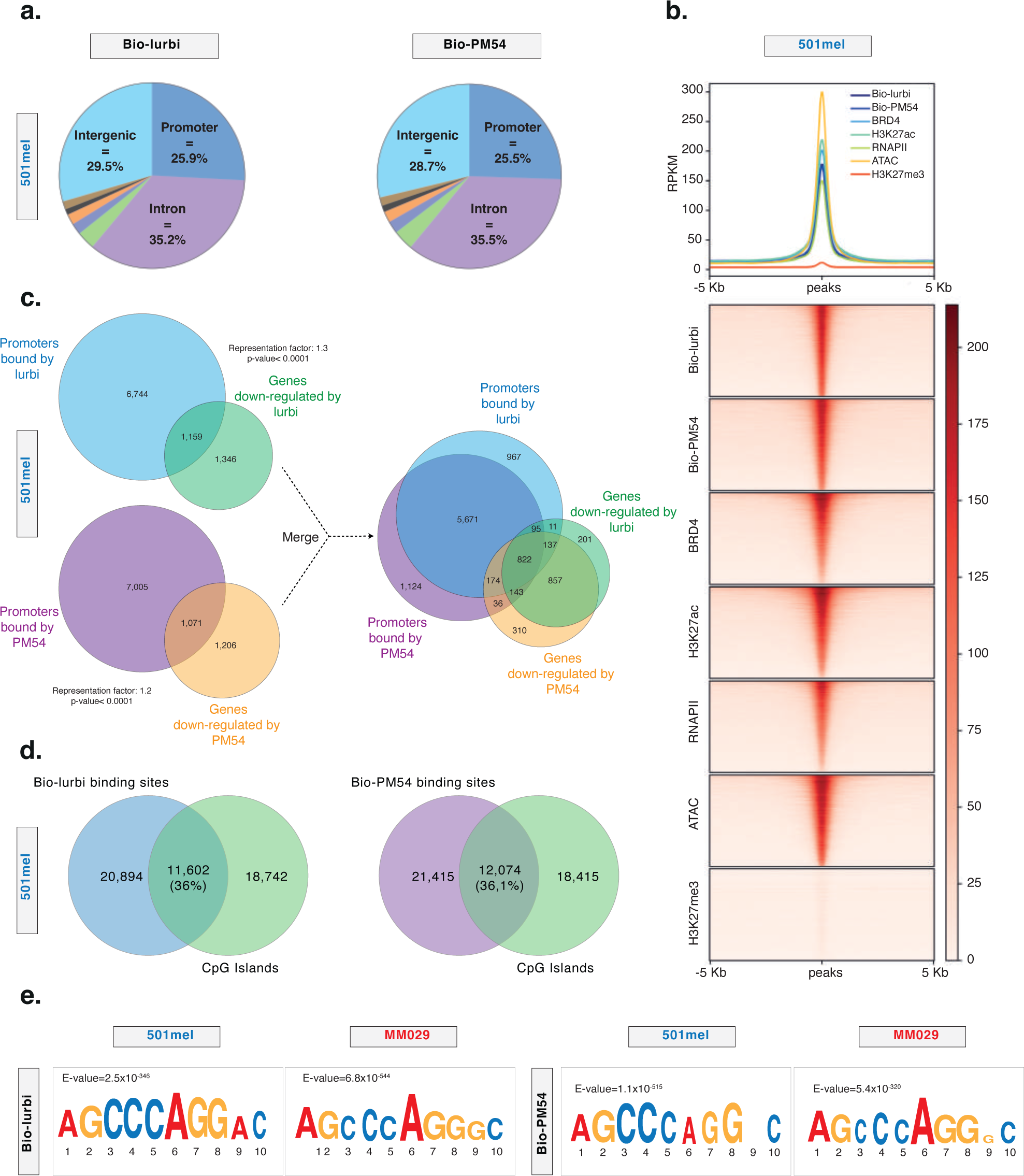
Synthetic ecteinascidins bind to promoter and intronic genomic regions. **a.** Pie chart showing the distribution of annotated peaks (in percentages) for Bio-lurbi (top) and Bio-PM54 (bottom) all over the genome (hg19) in 501mel cells. **b. Upper panel;** Metaplot distribution of Bio-lurbi, Bio-PM54, BRD4, RNAPII, H3K27ac, H3K27me3 enrichment and ATAC-Seq signals in a +/−5kb window around the occupied DNA binding sites of Bio-lurbi in differentiated 501mel cells. **Lower panel**; Heatmap profiles representing the read density clusterings obtained with seqMINER for the DNA-occupied sites of Bio-lurbi in differentiated 501mel cells relative to Bio-PM54, BRD4, RNAPII, H3K27ac, H3K27me3 enrichments and ATAC-Seq signals. Peak order is determined by Bio-lurbi and identical for all clusterings. **c. Left panel;** Venn diagram between promoters bound by Bio-lurbi or Bio-PM54 and genes down-regulated by lurbinectedin or PM54 in 501mel cells. **Right panel**; the two Venn diagrams were merged. Representation factor and hypergeometric p-value are indicated. **d.** Venn diagram between Bio-lurbi (top) and Bio-PM54 (bottom) binding sites in differentiated 501mel cells and human CpG Islands. **e.** Results of MEME-ChIP analysis on all lurbinectedin (top) and PM54 (bottom) occupied sites in either differentiated 501mel (left) or undifferentiated MM029 (right). Shown sequence represents the top enriched motif. E-values are shown.

Furthermore, our genome-wide analysis indicated that binding sites of synthetic ecteinascidins exhibited substantial overlaps with CpG islands in both melanoma cell types (**Figure 7d and Supplemental Figure 12e**). Employing the MEME-ChIP analysis tool facilitated an unbiased examination of the occupied sites. We identified a consistent CG-rich motif of 8 base pairs (bp) (AGCCCAGG) to be highly enriched across the binding sites identified for both drugs and in both cell types (**Figure 7e**). These data underscore the preferential binding of synthetic ecteinascidins at transcriptionally active, CG-rich genomic regions in melanoma cells and identified an 8 bp CG-rich motif as a preferential binding site for synthetic ecteinascidins.

### SE-related promoters occupied by synthetic ecteinascidins can be classified into different subgroups

We subsequently integrated the chemical-mapping data and observed a robust overlap (∼80%) between the promoter regions bound by Bio-lurbi and those bound by Bio-PM54 in a given cell type (**Figure 8a, left panel)**. Notably, among the promoters bound by synthetic ecteinascidins, 2,456 demonstrated concurrent binding by the two drugs in both cell types (**Figure 8a, right panel**). This included promoters that regulate the expression of ubiquitous transcription factors/coactivators such as CDK7, CDK9 or CDK12 and the Mediator subunits MED1 or MED13 **(Supplemental Table 6)**. In these promoters, synthetic ecteinascidins occupied CpG-rich sequences, which were typically strongly enriched in H3K27ac, RNAPII, and showed strong ATAC-seq signal, indicating actively transcribed genes (**Figure 8b**)

**Figure 8:**
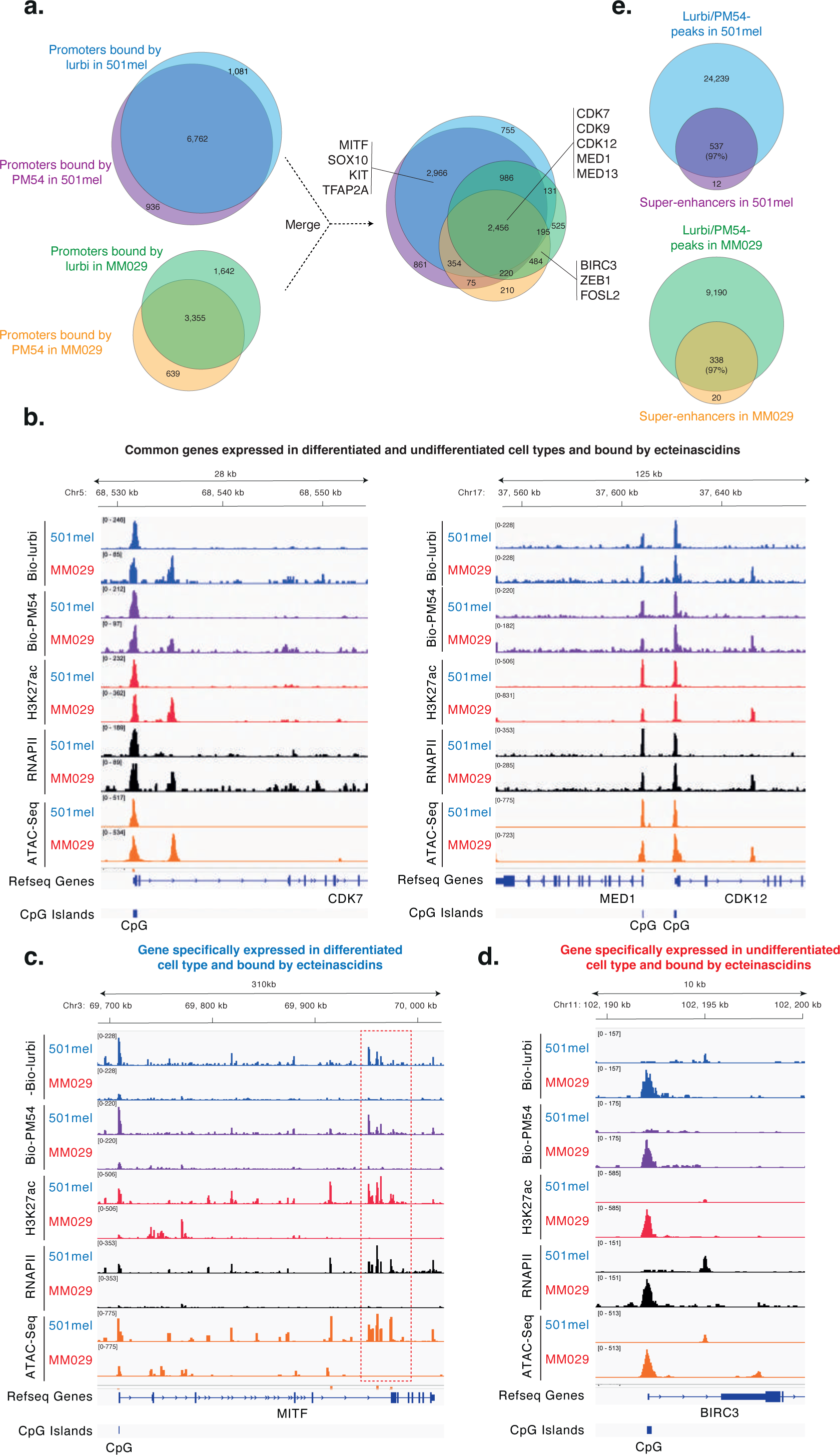
Synthetic ecteinascidins target the promoters of different subsets of genes. **a. Left panel;** Venn diagrams between promoters bound by Bio-lurbi and Bio-PM54 in 501mel (top) and MM029 (bottom) cells. **Right panel**; the two Venn diagrams were merged. **b.** Gene tracks of Bio-lurbi Bio-PM54, RNAPII, H3K27ac occupancy and ATAC-seq signals at *CDK7* (left) and *MED1/CDK12* (right) loci in 501mel or MM029 cells. These genes are expressed and bound by Bio-lurbi and Bio-PM54 in both 501mel and MM029 melanoma cells. **c.** Gene tracks of Bio-lurbi Bio-PM54, RNAPII, H3K27ac occupancy and ATAC-seq signals at the *MITF* locus in 501mel or MM029 cells. This gene is only expressed and bound by Bio-Lurbi and Bio-PM54 in 501mel cells. The red square indicates the SE regulating the expression of *MITF*. **d.** Gene tracks of Bio-lurbi Bio-PM54, RNAPII, H3K27ac occupancy and ATAC-seq signals at the *BIRC3* locus in 501mel or MM029 cells. This gene is only expressed and bound by Bio-lurbi and Bio-PM54 in MM029 melanoma cells. **e. Upper panel;** Venn diagrams between all genomic bindings sites commonly bound by Bio-lurbi and Bio-PM54 and *bona fide* super-enhancers identified in 501mel cells (top)**. Lower panel;** Venn diagrams between all genomic bindings sites commonly bound by Bio-lurbi and Bio-PM54 and *bona fide* super-enhancers identified in MM029 cells.

Apart from the commonality in drug-bound promoters depicted above, each melanoma cell type also exhibited a distinct pattern of binding associated with its specific cellular phenotype. Indeed, 2,966 and 484 promoters demonstrated exclusive binding in differentiated and undifferentiated cells, respectively. For instance, synthetic ecteinascidins bound to the promoter of the lineage-specific master transcription factor MITF only in differentiated cells, where it is highly expressed (**Figure 8c**). Conversely, the promoter of BIRC3, an inhibitor of apoptosis expressed only in undifferentiated melanoma cells, was occupied by synthetic ecteinascidins in undifferentiated but not in differentiated cells (**Figure 8d**).

When examining the deposition of synthetic ecteinascidins along the gene encoding MITF, drug-binding to its SE was also observed (**Figure 8c**). Similarly, the SE regulating the expression of FOSL2, a lineage-specific master transcription factor expressed in undifferentiated melanoma cells, was also occupied by synthetic ecteinascidins (**Supplemental Figure 13**). In agreement, almost all (∼95%) of the *bona fide* SEs identified by ROSE in differentiated and undifferentiated cells were directly bound by synthetic ecteinascidins (**Figure 8e**). Collectively, these findings suggest that synthetic ecteinascidins impact SE-mediated oncogenic transcription by binding to the promoters of ubiquitous transcription factors/coactivators enriched at SEs, together with the promoters of lineage-specific master transcription factors and potentially by directly targeting the SEs driving oncogenic expression.

### Synthetic ecteinascidins induce two waves of transcription inhibition in SKCM and SCLC

The data above suggest that the inhibition of ubiquitous transcription factors/coactivators may precede that of the SE-dependent genes. To test this hypothesis, we conducted kinetic analyses, revealing that transcription factors/coactivators were downregulated before SE-dependent oncogenes in SKCM cells (**Supplemental Figures 14a-d**). We explored whether a similar mechanism could occur in SCLC and NCSLC. Consistently, 351 genes were bound by synthetic ecteinascidins in SKCM and were commonly down-regulated in both SKCM, SCLC and NCSLC (**Supplemental Figure 15a**). Among these genes, the ubiquitous transcription factors/coactivators CDK7, CDK13, EP300, INO80 and the mediator subunit MED13 emerged. We observed that these genes were down-regulated very early in SCLC cells upon treatments with synthetic ecteinascidins, presumably leading to the inhibition of SCLC-specific SE-dependent oncogenic transcription (**Supplemental Figures 15b-c**). Overall, these data suggest that a first wave of inhibition affects transcription factors/coactivators in SKCM and SCLC treated with synthetic ecteinascidins, which then triggers the decommission of SEs and the inhibition of oncogenic expression.

## Discussion

Our working hypothesis posited that all types of melanoma cells, regardless of their specific phenotype and mutational statuses, would remain highly susceptible to the disruption of oncogene expression due to their inherent cancer-associated trait of transcriptional addiction ^30, 59, 35^. Melanoma cells display very high degrees of mutational burdens compared to other types of cancers, potentially resulting in proportional dysregulation of gene expression patterns. Moreover, the well-documented cell-state plasticity of melanoma cells underscores a robust reliance on tightly regulated oncogenic gene expression programs.

Comparative analyses were conducted to evaluate the impact of three synthetic ecteinascidins in relation to the clinically utilized MAPKi agents, namely vemurafenib, trametinib and dabrafenib. Notably, undifferentiated melanoma cells displaying inherent resistance to MAPKi, as well as *in vitro* engineered hyperpigmented cells with acquired MAPKi resistance ^47^, exhibited comparable sensitivity to the three synthetic ecteinascidins at concentrations within the low nanomolar range. *In vivo*, we observed potent decreases in mitotic indexes and increases in cell death and/or overall survival in four different melanoma CDX models, including MAPKi-resistant CDXs.

Our results shed light on the mechanisms of action of synthetic ecteinascidins, elucidating their common features, but also revealing some notable differential molecular effects. Low nanomolar doses of synthetic ecteinascidins commonly decreased proliferation and invasive capacities of melanoma cells, while inducing apoptosis and blocking the cell cycle in the S phase. We suspected the observed cellular effects to be at least partly due to DNA damage response signalling. Thus, we checked RNAPII degradation status and the induction of γH2AX and p-ATM. While the three compounds rapidly induced γH2AX in melanoma cells, a phenomenon not observed for MAPKi, marked differences were somewhat surprisingly observed between lurbinectedin and ecubectedin/PM54 treatments in some cellular models. In differentiated cells for example, RNAPII degradation and γH2AX were pronounced when treated with ecubectedin or PM54, and almost absent when cells were treated with lurbinectedin. These data highlight potential differences in efficacy and intracellular pharmacodynamics between the three compounds, which remain to be further studied. However, our data seem to argue that the degradation of RNAPII is not crucial for the cytotoxicity by synthetic ecteinascidins in melanoma cells.

Synchronously with the appearance of γH2AX, drug treatments also led to the important disruption of oncogene expression. Importantly, the transcriptional effects of the compounds seemed to exhibit a high degree of specificity for distinctly overexpressed oncogenes depending on the melanoma cell state. As such, while the expression of housekeeping genes was not affected in 2- or 3-D conditions by short-term drug treatments, lineage-specific drivers of proliferation such as MITF, SOX10 or PAX3 were strongly inhibited specifically in differentiated cells. In undifferentiated cells however, different genes were affected, such as the key regulators AXL or EGFR, the anti-apoptotic protein BIRC3 or the cell-type master transcription factors FOSL2 and TEAD4. These observations reveal arguably the most interesting feature of these novel compounds; synthetic ecteinascidins selectively bind to highly transcribed genomic regions and seem to specifically inhibit the distinct transcription programs on which a given cancer cell subpopulation depends on. Thus, the efficacy of synthetic ecteinascidins does not depend on the phenotypic nature of the melanoma cell, a feature that differentiates these drugs from conventional MAPKi therapies and immunotherapy ^60^. Therefore, our findings underscore the potential clinical benefit of using these novel compounds as a second-line treatment after MAPKi/immunotherapy relapse.

Mechanistically, our results highlight a multifaceted mechanism of action by which synthetic ecteinascidins impede oncogenic transcription. Synthetic ecteinascidins bind to promoters of genes encoding ubiquitous transcription factors/coactivators usually strongly enriched at SEs, leading to their rapid inhibition. The above-mentioned effect is most likely potentiated by the fact that genes encoding for lineage-specific master transcription factors such as MITF or FOSL2 are also heavily bound by synthetic ecteinascidins in both differentiated and undifferentiated melanoma cells. These regulators are known to bind SEs to form autoregulatory loops forming the core transcriptional regulatory circuitries of melanoma cells. The disruption of these oncogenic expression loops, added to the fact that synthetic ecteinascidins also seem to strongly bind SEs themselves, albeit with uncertain biological consequences, potentially further ensures the strong inhibition of SE-driven oncogenic transcription.

Delving deeper into the transcriptional effects elicited by the three compounds, we observed that while the gene expression changes elicited by lurbinectedin and ecubectedin greatly overlapped, the transcriptional effects of PM54 significantly diverged. Notably, PM54 treatments deregulated fewer genes than lurbinectedin or ecubectedin, while eliciting the same cytostatic and cytotoxic effects, thus representing potentially a clinical benefit. Although the exact mechanism explaining this difference is unknown, it may be related to the fact that the moiety that is modified in PM54 *vs.* lurbinectedin is located in the area of the molecule described as interacting with DNA binding proteins/transcription factors ^61^. Such a differential interaction between the drug and transcription factors might cause less systemic gene expression disruptions and thus unwanted secondary effects while still potently targeting the promoters of transcription factors/coactivators and lineage-specific master transcription factors, leading to cancer cell death. Consequently, Phase I clinical trials for PM54 in advanced solid tumors, including melanoma, were initiated (ClinicalTrials.gov Identifier: NCT05841563).

Collectively, our data allow for a comprehensive overview of the cellular and molecular effects of a potential novel therapeutic approach to melanoma, based on the dual mechanism of action of DNA damage induction and SE-dependent oncogenic inhibition. The current study further sheds light onto the intricacies of gene expression dependencies of different melanoma cell subpopulations and their molecular reactions towards transcriptional disruptions. While this important preclinical work might legitimize the clinical testing of synthetic ecteinascidins, it also highlights the potential benefits of further exploring the effects of additional structural analogues.

## Materials and Methods

### Resource availability

#### Lead contact

Further information and requests for resources and reagents should be directed to and will be fulfilled by the Lead Contact, Frédéric Coin

#### Extended resource table

An extended resource table with antibodies, oligonucleotide sequences, chemicals and reagents used in this work is provided in **Supplemental Table 7**.

#### Data and code availability

Next generation sequencing raw and processed data have been deposited at GEO: Accession numbers of these data are listed in the **Supplemental Table 7**. This paper analyzes existing, publicly available data. These accession numbers for the datasets are listed in **Supplemental Table 7**.

### Cell culture and treatment

Cells were grown at 37°C in 5% CO2 (10% for Hermes 3A) and were regularly checked for mycoplasma contamination. MM patient-derived short-term melanoma cultures (MM011, MM074, MM117, MM029, MM047, MM099) were grown in HAM-F10 (Gibco, Invitrogen) supplemented with 10% Fetal Calf Serum (FCS), 25 mM HEPES, 5,2 mM GLUTAMAX and penicillin–streptomycin. Melanoma cell lines 501mel and SKmel28 were grown in RPMI w/o HEPES (Gibco, Invitrogen) supplemented with 10% FCS and gentamycin. Vemurafenib-resistant cells (501mel^VemuR^ and MM074^VemuR^) were additionally supplemented with 1,5μM of vemurafenib. Melanoma IGR cell lines (IGR37 and IGR39) were grown in RPMI w/o HEPES (Gibco, Invitrogen) supplemented with 15% FCS and gentamycin. Immortalized melanocytes Hermes-3A were grown in RPMI w/o HEPES supplemented with 10% FCS, penicillin-streptomycin, 200nM TPA (Sigma Aldrich), 200p.m. Cholera Toxin (Sigma Aldrich), 10 ng/mL hSCF (Life Technologies), 10nM EDN-1 (Sigma Aldrich) and 2mM Glutamine (Invitrogen). Small cell lung cancer cell line DMS53 was grown in Waymouth’s MB medium (Gibco, Invitrogen), supplemented with 10% FCS and gentamycin. 501mel, SKmel28, IGR and DMS53 cells were purchased from ATCC, MM and Hermes-3A cells were obtained from collaborators. Vemurafenib (PLX4032), trametinib (GSK1120212) and dabrafenib (GSK2118436) were purchased from Selleckchem. Lurbinectedin (PM1183), ecubectedin (PM14), and PM54 were obtained from PharmaMar S.A. Recombinant Human IFN-γ was obtained from Peprotech (300-02).

### Protein extraction and Western Blotting

For whole cell extracts, cells were rinsed once with cold PBS, before pelleting and resuspension in LSDB 0.5M buffer (500 mM KCl, 50 mM Tris pH 7.9, 20% glycerol, 1% NP-40, 1mM DTT and protease inhibitor cocktail). Afterwards, cells were fully disrupted with 3 cycles of heat shock (liquid nitrogen followed by 37°C water bath). Then, samples were centrifugated for 15 minutes at 14,000rpm to remove cell debris. Lysates were subjected to SDS–polyacrylamide gel electrophoresis (SDS-PAGE) and proteins were transferred onto a nitrocellulose membrane. Membranes were incubated overnight 4 °C with primary antibodies in PBS+ 5% milk powder + 0.01% Tween-20. The membranes were then incubated with HRP-conjugated secondary antibody (Jackson ImmunoResearch) for 1 hour at room temperature and visualized using the ECL detection system (GE Healthcare).

### IC50 estimation

Cells were seeded at 5×10^3^ cells/well in 96-well plates and treated with increasing concentrations of vemurafenib, dabrafenib, trametinib, lurbinectedin, ecubectedin, or PM54. After 72 hours of incubation, cells were treated with PrestoBlue reagent (ThermoFisher) according to the manufacturer’s instructions. The absorbance per well was measured with a CellInsight CX5 microplate reader (ThermoFisher). Determination of IC50 values was performed by nonlinear curve fitting using the Prism9 statistical software (GraphPad). To assess the effect of IFNγ on drug sensitivities, cells were pre-treated with IFNγ (20 ng/mL) for 24 hours, before being treated as mentioned above, while maintaining IFNγ (20 ng/mL) in the medium.

### Clonogenicity Assay

Cells were drug-treated at IC50 concentrations during 48 hours before seeding 1×10^3^ or 2×10^3^ cells in 6-well plates without drugs, where they grew for 10 days to allow for colony formation. Afterwards, cells were fixed for 10min with 4% Formaldehyde solution, washed once with PBS and stained with Crystal Violet solution 0.2% for 15 minutes. The wells were finally washed twice with deionized water, air dried, scanned and analyzed with Fiji software to count the number of colonies.

### Cell proliferation, apoptosis, and cell cycle analysis by Flow Cytometry

2×10^6^ cells were seeded in 6 well plates and were incubated 24 hours later with 1uM of CellTrace Violet reagent (ThermoFisher) according to the manufacturer’s instructions, immediately before rinsing and drug treatment at IC50 concentrations. After 48 hours of incubation, cells were rinsed and incubated with AnnexinV-APC (BD Biosciences). Cell proliferation and apoptosis were detected on a BD LSRFortessaTM Flow Cytometer. Data were analysed with FlowJo software. To define slow proliferating or apoptotic cells, we proceeded as follows: We considered that slow proliferating cells represented the 30% of cells with the highest concentration of CellTrace Violet signal in the DMSO control. We then calculated the % of cells that had a signal greater than or equal to this value with drug treatment. For apoptotic cells, we considered the 20% of cells with the highest signal of AnnexinV-APC in the DMSO control.

For cell cycle analysis, 2×10^6^ cells were seeded in 6 well plates. After 72 hours of drug treatments at IC50 concentrations, cells were pelleted and fixed with 70% ethanol for 1h at 4°C. After 2 washes with cold PBS, cells were incubated with RNAseA and PI for 1 hour in the dark, before being analyzed on a BD LSRFortessaTM Flow Cytometer. Data were analysed with FlowJo software.

For apoptosis assays with 3D-grown melanoma cells, TrypLe Select 10x reagent (Gibco) was used to dissociate melanospheres to obtain single-cell suspensions. These cells were incubated with AnnexinV-APC (Biolegend) and Propidium Iodide (PI, Biolegend). With bivariant dot plots, we distinguished between viable (AnnexinV− / PI−), early apoptotic (AnnexinV+ / PI−), late apoptotic (AnnexinV+ / PI+) and necrotic cells (AnnexinV− / PI+).

### Boyden Chamber Invasion Assay

2×10^6^ cells were seeded inside Boyden Chamber inserts (Fisher Scientific) with 4% Matrigel (Corning) and covered with serum-free media. The inserts were placed in 24 well plates filled with complete medium. After 24 hours, the inserts were fixed for 10min with 4% Formaldehyde solution, washed once with PBS and stained with Crystal Violet solution 0.2% for 15min. The wells were finally washed twice with deionized water, air dried, and photos were collected using an EVOS xl Core microscope. The pictures were analyzed with Fiji to assess the area of occupancy of the cells.

### Wound-healing assay

Confluent melanoma cell monolayers in 6-well plates were scratched with a 20-µL pipette tip to create uniform, cell-free wounds. Fresh medium without FCS (to mitigate proliferation), with or without drugs, was added. At 0, 24, and 48 hours, photomicrographs of the wounds were taken under an inverted microscope. The wound areas were then quantified using Fiji software.

### Melanosphere formation and viability assay

5×10^4^ cells were seeded in ultra-low attachment hydrogel-layered 96 well plates (Corning 7007) in KO DMEM medium supplemented with 20% KSR, AANE, 2 mM Glutamax, Penicillin/Streptomycin and 100 uM Beta-mercaptoethanol. To allow for melanosphere formation, cells were left to grow for 4 days before drug treatment.

To analyze melanosphere viability after drug treatment, cells were treated with CellTiterGlo reagent (Promega) according to the manufacturer’s instructions. Luminescence signals were measured with a Centro XS LB 960 microplate reader (Berthold).

### RNA Extraction and RT-qPCR

Total RNA isolation was performed according to the manufacture protocol with NucleoSpin RNA Plus kit (Macherey-Nagel). RNA was retrotranscribed with Reverse Transcriptase Superscript IV (Invitrogen), qPCR was performed with SYBR Green (Roche) and on a LightCycler 480 (Roche). Target gene expression was normalized using 18S as reference gene.

### Bulk RNA-Sequencing and analysis

Library preparation was performed at the GenomEast platform at the Institute of Genetics and Molecular and Cellular Biology using TruSeq Stranded Total RNA Reference Guide - PN 1000000040499. Total RNA-Seq libraries were generated from 700 ng of total RNA using TruSeq Stranded Total RNA Library Prep Gold kit and TruSeq RNA Single Indexes kits A and B (Illumina, San Diego, USA), according to manufacturer’s instructions. Briefly, cytoplasmic and mitochondrial ribosomal RNA (rRNA) was removed using biotinylated, target-specific oligos combined with Ribo-Zero rRNA removal beads. Following purification, the depleted RNA was fragmented into small pieces using divalent cations at 94oC for 8 minutes. Cleaved RNA fragments were then copied into first strand cDNA using reverse transcriptase and random primers followed by second strand cDNA synthesis using DNA Polymerase I and RNase H. Strand specificity was achieved by replacing dTTP with dUTP during second strand synthesis. The double stranded cDNA fragments were blunted using T4 DNA polymerase, Klenow DNA polymerase and T4 PNK. A single ‘A’ nucleotide was added to the 3’ ends of the blunt DNA fragments using a Klenow fragment (3’ to 5’exo minus) enzyme. The cDNA fragments were ligated to double stranded adapters using T4 DNA Ligase. The ligated products were enriched by PCR amplification. Surplus PCR primers were further removed by purification using AMPure XP beads (Beckman-Coulter, Villepinte, France) and the final cDNA libraries were checked for quality and quantified using capillary electrophoresis. Libraries were sequenced on an Illumina HiSeq 4000 sequencer as single read 50 base reads. Image analysis and base calling were performed using RTA version 2.7.7 and bcl2fastq version 2.20.0.422.

Reads were preprocessed to remove adapter and low-quality sequences (Phred quality score below 20). After this preprocessing, reads shorter than 40 bases were discarded for further analysis. These preprocessing steps were performed using cutadapt version 1.10. Reads were mapped to rRNA sequences using bowtie version 2.2.8 and reads mapping to rRNA sequences were removed for further analysis. Reads were mapped onto the hg19 assembly of Homo sapiens genome using STAR version 2.5.3a. Gene expression quantification was performed from uniquely aligned reads using htseq-count version 0.6.1p1, with annotations from Ensembl version 75 and “union” mode. Only non-ambiguously assigned reads have been retained for further analyses. Read counts have been normalized across samples with the median-of-ratios method proposed by Anders and Huber ^45^ to make these counts comparable between samples. Comparisons of interest were performed using the Wald test for differential expression ^62^ and implemented in the Bioconductor package DESeq2 version 1.16.1. Genes with high Cook’s distance were filtered out and independent filtering based on the mean of normalized counts was performed. P-values were adjusted for multiple testing using the Benjamini and Hochberg method ^63^. Deregulated genes were defined as genes with log2(Fold change) > 1 or < - 1 and adjusted P-value < 0.05.

Volcano plots were generated using the Prism9 statistical software (GraphPad). Heatmaps were generated using Morpheus (https://software.broadinstitute.org/morpheus). Venn diagrams were generated using DeepVenn (http://www.deepvenn.com/) and representation factors and hypergeometric P-values were determined using Graeber lab software (https://systems.crump.ucla.edu/hypergeometric/). Gene Ontology Analysis was performed using ShinyGO ^64^.

### In vitro Immunofluorescence Assays

After PBS-rinsing, cells grown on coverslips were fixed with 4% PFA for 15 min. Cells were then permeabilized with PBS and 0.1% Triton X-100. Blocking was done with 10% BSA. Primary antibodies were incubated overnight at 4°C, after which cells were stained for 1 hour at room temperature with AlexaFluor-conjugated secondary antibodies diluted in PBS+10% FCS (Life technologies) and stained with DAPI. For γH2AX quantifications, image acquisition was performed on a DFC7000T widefield microscope (Leica) and γH2AX signals were assessed for each DAPI-positive area using the Fiji software. For BRD4 and MED1 foci quantifications, image acquisition was performed on a TCS SP5 inverted confocal microscope (Leica), and foci were counted using the Cell Counter plugin of the Fiji software.

### Immunofluorescence on tumor sections

Tumors were grown as mentioned above and were extracted after 24 hours following a single dose of placebo treatment or 1.2 mg/kg of Ecubectedin or PM54. In parallel, untreated tumors were extracted. The tumors were fixed in 10% formalin and embedded in paraffin for histology. Slides prepared from 5μm-thick paraffin sections were processed for antigen retrieval in 10 mM sodium citrate buffer (PH = 6.0) for 45 minutes at 95°C in a water bath. The slides were cooled down at room temperature (RT) for 15 minutes. They were rinsed in PBS and then incubated in a humidified chamber for 16 hours at 4 °C, with the primary antibodies diluted in PBS containing 0.1% (v/v) Tween 20 (PBST) to detect mitotic (pHH3-positive) and apoptotic (cleaved caspase 3-positive) cells. After rinsing in PBST, detection of the bound primary antibodies was performed for 1 hour at room temperature in a humidified chamber using 555-conjugated secondary rabbit IgG antibody. The sections were then counterstained with DAPI to label nuclei. Stained sections were digitalized using a slide scanner (Nanozoomer 2.0-HT, Hamamatsu) and analyzed with the corresponding ND.view2 software.

Large 8-Bits digital scanned images of tumors stained for nuclei (10 000 to 30 000 nuclei per section) and pHH3 or cleaved caspase 3 were processed through an inhouse python (v3.8) algorithm to quantify positive cells. Basically, blue channels were proposed to a Cellpose2 model (deep learning model backboned by pytorch process) to segment nuclei. Subsequently, nuclei were analyzed for specific signals. For pHH3, a nucleus was considered positive if total pixels above 50 in intensity value exceeds 20% of nuclei surface (in 8 Bits image values range from 0 [no signal] to 255). Hence, we ensured that we did not consider unspecific background signals or insignificantly bright signals. The same procedure was applied to Caspase3 with pixel value set to 50 and minimal covered surface set to 30%. For each image, a ratio of positive cells/total nuclei was returned as the experimental variable. Statistics were produced using python’s pingouin library (v0.5.3) with two-way ANOVA and post hoc tests being built-in functions.

### Xenograft models

4- to 6-week-old NSG or athymic nude female mice were subcutaneously implanted into their right flank with human melanoma cell suspensions (LOX-IMVI, WM-266-4, 501mel, or 501mel^VemuR^). When tumors began to develop, these were measured 2-3 times per week. Tumor volume was calculated with the equation (a x b^2^)/2, where “a” and “b” referred to the longest and shortest diameters, respectively. When tumors reached a size of 150 mm3, tumor bearing animals (N = 8/group) were treated with Placebo (saline solution) or ecubectedin or PM54 at 1.2 mg/kg weekly. Tumor volume and animal body weights were measured 2-3 times per week, starting from the first day of treatment. The median was determined for tumor volume/size on each measurement day. Treatment tolerability was assessed by monitoring body weight evolution, clinical signs of systemic toxicity, as well as evidences of local damage in the injection site. Treatments which produced >20% lethality and/or 20% net body weight loss were considered toxic. Furthermore, animals were euthanized when their tumors reached ca. 1500 mm3 and/or severe necrosis was seen. Differences on antitumor effect were evaluated by comparing tumor volume data as well as median survival time from the placebo treated group with Ecubectedin or PM54 treated groups. For this, a two-tailed Mann-Whitney U test was used.

### Chemical-mapping and Cut&Tag

501mel and MM029 cells were seeded and grown to sub-confluency in 15-cm plates before treatment for 8 hours with DMSO, biotinylated lurbinectedin (Bio-lurbi) or biotinylated PM54 (Bio-PM54) at a concentration equivalent to 10xIC_50_ (15nM for Bio-lurbi and 30nM for Bio-PM54). Chemical-mapping and CUT&TAG were then performed using the Active Motif CUT&Tag-IT assay kit (53160, 53165), following the manufacturer’s instructions. Briefly, 5×10^5^ cells per condition were collected and washed twice before being bound to Concanavalin A beads and then incubated overnight at 4°C with primary antibodies (1:50 dilutions). The following day, the corresponding guinea pig Anti-rabbit or rabbit Anti-mouse secondary antibodies were used at a 1:100 dilution in digitonin buffer and incubated at room temperature for 1 hour. Subsequently, the CUT&Tag-IT Assembled pA-Tn5 Transposomes were incubated at room temperature for 1 hour, and cells were resuspended in Tagmentation buffer and incubated at 37°C for 1 hour. The Tagmentation process was then stopped by adding EDTA and SDS. Protein digestion was performed by adding Proteinase K (10 mg/mL) and incubating at 55°C for 1 hour. The DNA was retrieved with DNA purification columns provided by the manufacturer and was then subjected to library preparation and PCR amplification and purified by 2 successive washes with SPRI beads. Libraries were sequenced on an Illumina NextSeq 2000 sequencer as paired-end 50 base reads. Image analysis and base calling were performed using RTA version 2.7.7 and BCL Convert version 3.8.4. The adapter sequence: CTGTCTCTTATA has been trimmed with cutadapt 1.18 with option: -a CTGTCTCTTATA -A CTGTCTCTTATA -m 5 -e 0.1 and Bowtie2 ^65^ parameter: -N 1 -X 1000, was used for mapping to the human genome (hg19). After the mapping, reads overlapping with ENCODE blacklist V2 were filtered. Each de-duplicated read was extended to its fragment size. Tracks were normalized with RPKM method. Peak calling was performed using Macs2 ^66^ 2.2.7.1 in BEDPE and narrow mode. narrowPeaks from biological triplicate samples were then merged to a single master peak set. BEDtools ^67^ was used to calculate the read coverage for each peak and for each sample. Peaks were annotated using Homer ^68^ software with ucsc 6.4 gene annotation. Bigwig tracks were generated using bamCoverage from deepTools 3.5.4 ^69^. The differential analysis was performed using DESeq2 ^70^. Peak correlation analysis was performed using DiffBind ^71^ r package. Heatmap and average profile analyses were performed using seqMINER ^72^ and deepTools. Motif analysis was performed using MEME-ChIP ^73^ with JASPAR 2020 core vertebrates motif collection. For Super-Enhancer calling, ROSE algorithm version 0.1 (http://younglab.wi.mit.edu/super_enhancer_code.html) was applied with default parameters (stitch distance = 12500, ^74, 75^) using the BRD4 or H3K27ac peaks identified by MACS2 with the Cut&Tag experiments. TSS regions (Refseq TSS ±1000bp) were excluded.

### ATAC-Seq

501mel and MM029 cells were seeded and grown to sub-confluency in 15-cm plates, and ATAC-Seq was then performed using the Active Motif ATAC-Seq Kit (53150), following the manufacturer’s instructions. Briefly, 1×10^5^ nuclei were isolated by adding 100 μL ice cold ATAC-lysis buffer to the cell pellet. After centrifugation (500 g, 10 minutes at 4°C), cells were washed and incubated with the tagmentation master mix in a shaking heat block at 37°C/800 rpm for 30 minutes. Obtained DNA was taken up in DNA purification buffer, purified using the contained DNA purification columns, amplified for 10 cycles using indexed primers, and size-selected using SPRI beads. Libraries were sequenced on an Illumina NextSeq 2000 sequencer as paired-end 50 base reads. Image analysis and base calling were performed using RTA version 2.7.7 and BCL Convert version 3.8.4. Samples were analyzed using the ENCODE ATACseq pipeline release v2.0.2 with hg19 assembly.

### ChIP-qPCR

501mel and MM029 cells were seeded and grown to sub-confluency in 15-cm plates. After drug treatments, cells were fixed with 0.4% PFA for 10 min and quenched with 2 M Glycin pH 8. Cells pellets were lysed in 25 mM HEPES pH 7.8, 10 mM NaCl, 1.5 mM MgCl_2_, 0.5% NP-40, 1 mM DTT. Nuclei were resuspended in in 50 mM Hepes-KOH pH 7.8, 140 mM NaCl, 1 mM EDTA, 1% Triton X-100, 0.1 mM Na-deoxycholate, 0.1% SDS and sonicated at 4°C with a Q500 sonicator (Qsonica) to get DNA fragments between 100-500 bp. 50 µg of the sonicated chromatin was then diluted in Dilution buffer (1% Triton X-100, 2 mM EDTA, 20 mM Tris HCl pH 7.5, 150 mM NaCl) and incubated overnight at 4°C with 5 ug of respective antibodies. The antibody-chromatin complex was then captured with a mix of protein A and G Dynabeads (Invitrogen) for 2 hours at 4°C, and beads were then washed twice in Low Salt Washing Buffer (1% Triton, 2 mM EDTA, 20 mM Tris HCl pH 7.5, 150 mM NaCl, 0.1% SDS), High salt Washing Buffer (1% Triton, 2 mM EDTA, 20 mM Tris HCl pH 7.5, 500 mM NaCl, 0.1% SDS), and TE buffer (100 mM Tris HCl pH 7.5, 10 mM EDTA). Immunoprecipitated chromatin was subsequently eluted from beads in 1% SDS and 100mM NaHCO_3_ at 65°C for 30 minutes, and crosslinks were reversed by overnight incubation with Proteinese K (50µg/ml) at 65 °C. The DNA was finally purified with the QIAquick PCR Purification kit (QIAGEN), resuspended in 200 µL of water, and analyzed by qPCR. Quantification of ChIP DNA concentrations with qPCR was performed by calculating the percent of input for each ChIP sample, calculated as 2^(Ct_input -Ct_IP) × 100. Subsequently, the obtained percentage was normalized to the negative control IgG. Finally, the fold enrichment of the drug-treated samples over the DMSO-treated samples was calculated.

### Statistics and reproducibility

Experimental data was plotted and analyzed using either Excel (Microsoft) or GraphPad Prism (GraphPad Software Inc.). The number of samples and replicates are indicated in the respective figure legends.

## Author contributions

Conceptualization: FC

Methodology: MC, JO, ID, CC, FC

Investigation: MC, JO, MN, LS, PC, PB, CC, CM, AZ, CE, TKL, EC, PA

Analysis of data: TY, GD

Funding acquisition: FC, JME, CC

Supervision: FC, CC

Writing – original draft: FC

Writing – review & editing: ID, EC, JME, CC

## Acknowledgements

This study was supported by the Ligue contre le cancer (Equipe Labélisée 2022-2024), the Institut National du Cancer (INCa) (PLBIO23-008), the grant ANR-10-LABX-0030-INRT, a French State fund managed by the Agence Nationale de la Recherche under the frame program Investissements d’Avenir ANR-10-IDEX-0002-02. This work of the Interdisciplinary Thematic Institute IMCBio+, as part of the ITI 2021-2028 program of the University of Strasbourg, CNRS and Inserm, was supported by IdEx Unistra (ANR-10-IDEX-0002), and by SFRI-STRAT’US project (ANR-20-SFRI-0012) and EUR IMCBio (ANR-17-EURE-0023) under the framework of the France 2030 Program. Sequencing was performed by the IGBMC GenomEast platform, a member of the “France Génomique” consortium (ANR-10-INBS-0009).□ Histology and subsequent image analysis were performed by the pathology facility of the mouse clinic institute (ICS), a member of the Celphedia/Phenomin infrastructures (ANR-10-INBS-0007). We also thank the IGBMC Cell culture, Flow cytometry, Phenotypic analysis and cellular screening, and Photonic microscopy services. P.B is supported by the Ligue contre le Cancer. JME was sponsored by a Mount Jade Scholar Fellow from the MOST of Taiwan and a Yonglin Chair Professor grant of National Taiwan University. We thank Dr Hsiang-Hung Huang for having initiated the work on SCLC cells.

## Supplemental Tables and Figures

**Supplemental Table 1:** List of genes with their relative expression *vs.* DMSO treatment for several cancer cells following treatment with either lurbinectedin, ecubectedin or PM54.

**Supplemental Table 2:** List of genes displaying consistent down (757)- and up (110)-regulation across both MM074 and MM029 cells in response to all the three compounds (lurbinectedin, ecubectedin or PM54).

**Supplemental Table 3:** List of genes displaying consistent down-regulation across cells from SKCM, NSCLC and SCLC in response to lurbinectedin.

**Supplemental Table 4:** List of SEs identified by ROSE following Cut&Tag against H3K27ac or BRD4 in 501mel and MM029 is provided on page 1. A list of SE-dependent genes identified by ROSE in 501mel and MM029 is provided on page 2. A list of SE-dependent genes down-regulated in 501mel or MM029 cells following treatment with either lurbinectedin, ecubectedin or PM54 is provided on page 3.

**Supplemental Table 5:** The list of genome-wide binding sites for Bio-lurbinectedin and Bio-PM54 in 501mel and MM029 cells as determined by Chemical-mapping.

**Supplemental Table 6:** The list of promoters/TSS-bound genes for Bio-lurbinectedin and Bio-PM54 in 501mel and MM029 as determined by Chemical-mapping.

**Supplemental Table 7:** An extended resource table with antibodies, oligonucleotide sequences, chemicals and reagents used in this work.

**Supplemental Figure 1:**
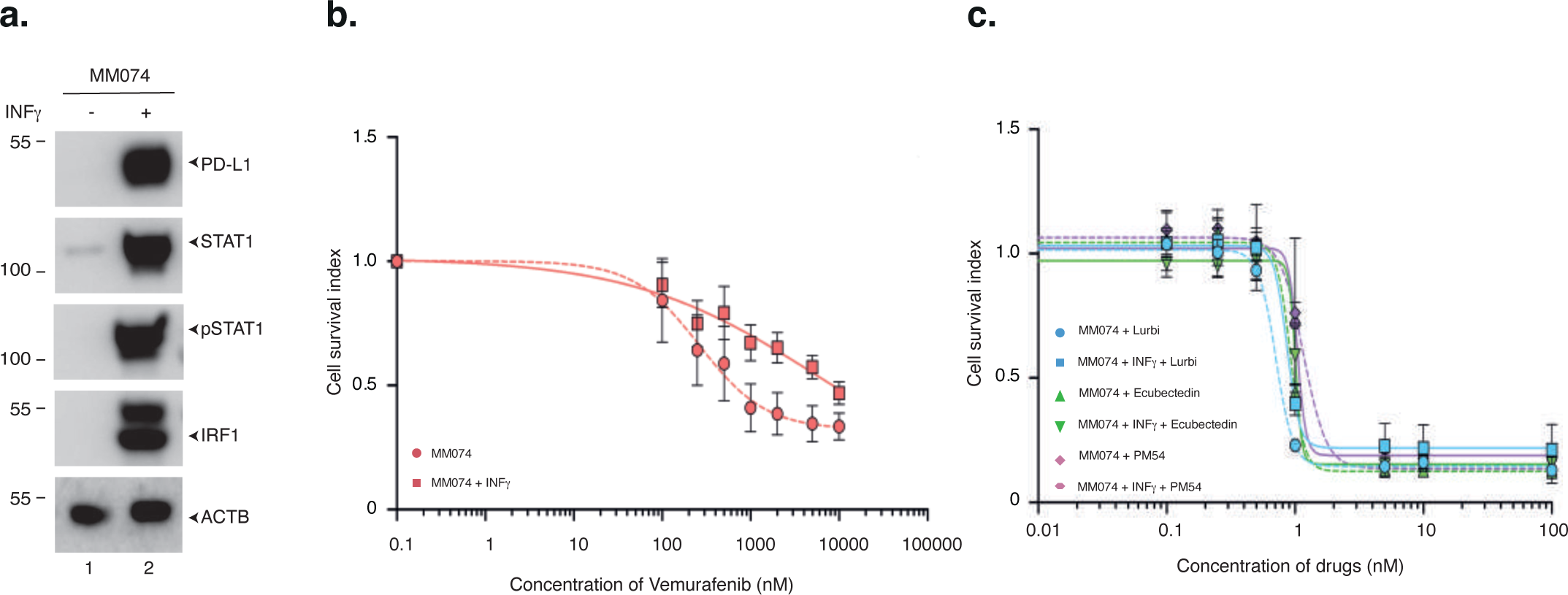
Pseudo-interferon-active melanoma cells are resistant to MAPKi but sensitive to synthetic ecteinascidins. **a.** Protein lysates from differentiated MM074 melanoma cells treated or not with INFγ (20ng/ml, 24 hours), were immuno-blotted for proteins as indicated. Molecular mass of the proteins is indicated (kDa). **b.** MM074 cells were pre-treated with INFγ (20ng/ml, 24 hours) and then with increasing doses of vemurafenib, in the presence of INFγ. Mean growth is shown relative to vehicle (H2O)-treated cells. Error bars indicate mean values +/− Standard Deviation (SD) for three biological triplicates. **c.** MM074 cells were treated with INFγ for (20ng/ml, 24 hours), and then with increasing concentrations of lurbinectedin, ecubectedin or PM54, in the presence of INFγ. Mean growth is shown relative to mock-treated cells. Error bars indicate mean values +/− Standard Deviation (SD) for three biological triplicates.

**Supplemental Figure 2:**
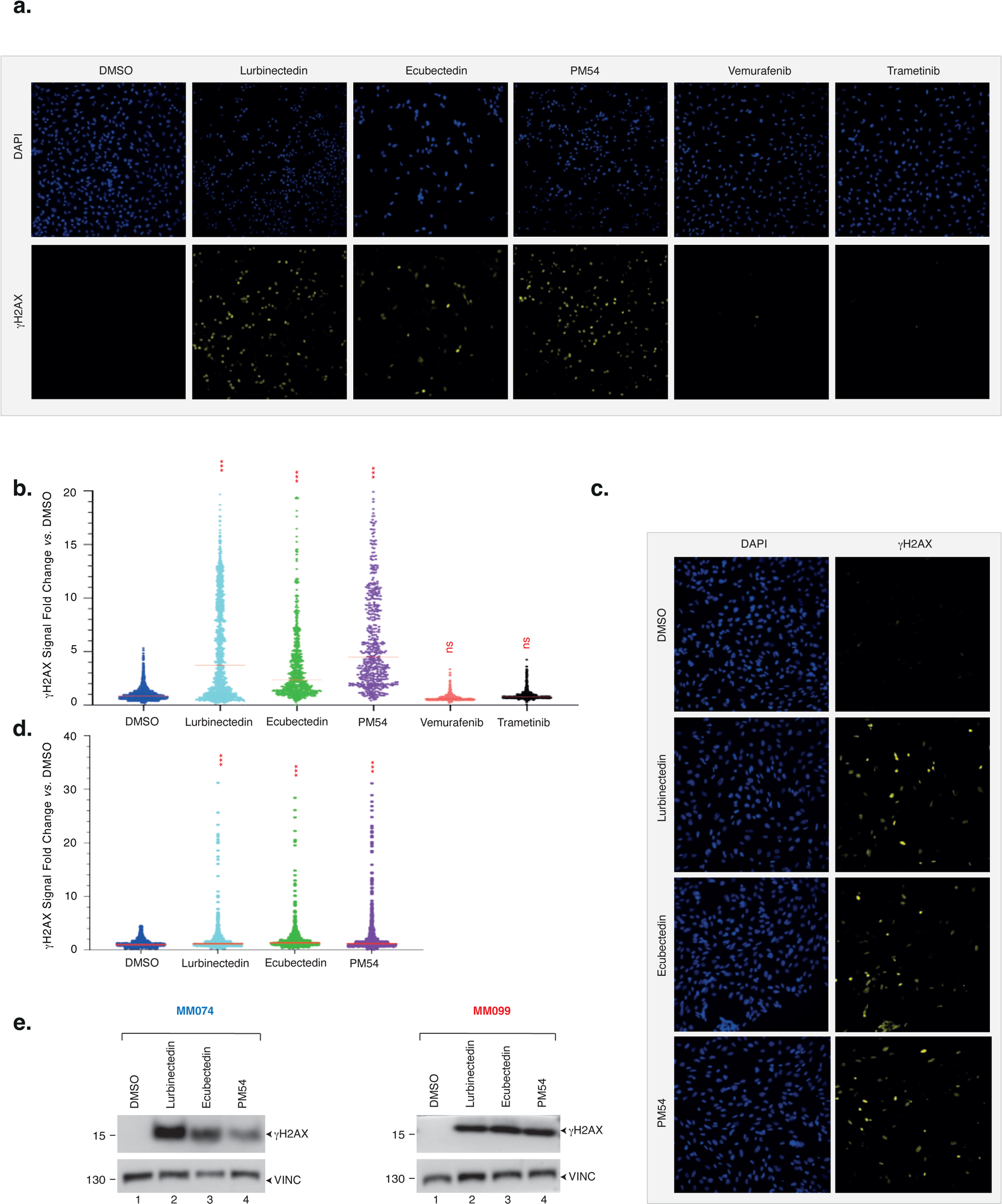
Synthetic ecteinascidins potently induce DSBs. **a-d.** Differentiated 501mel (**a, b**) or undifferentiated MM029 (**c, d**) cells were treated with indicated drugs (5xIC50 concentration, 24 hours) and γH2AX induction was assessed by immunofluorescence. Representative images are shown (**a, c**) as well as γH2AX signal quantification (n= at least 500 nuclei in three independent experiments). Bars indicate mean values (**b, d**). **e.** Protein lysates from differentiated MM074 or undifferentiated MM099 treated with either lurbinectedin, ecubectedin or PM54 (5xIC50 concentration, 24 hours), were immuno-blotted for proteins as indicated. Molecular mass of the proteins is indicated (kDa).

**Supplemental Figure 3:**
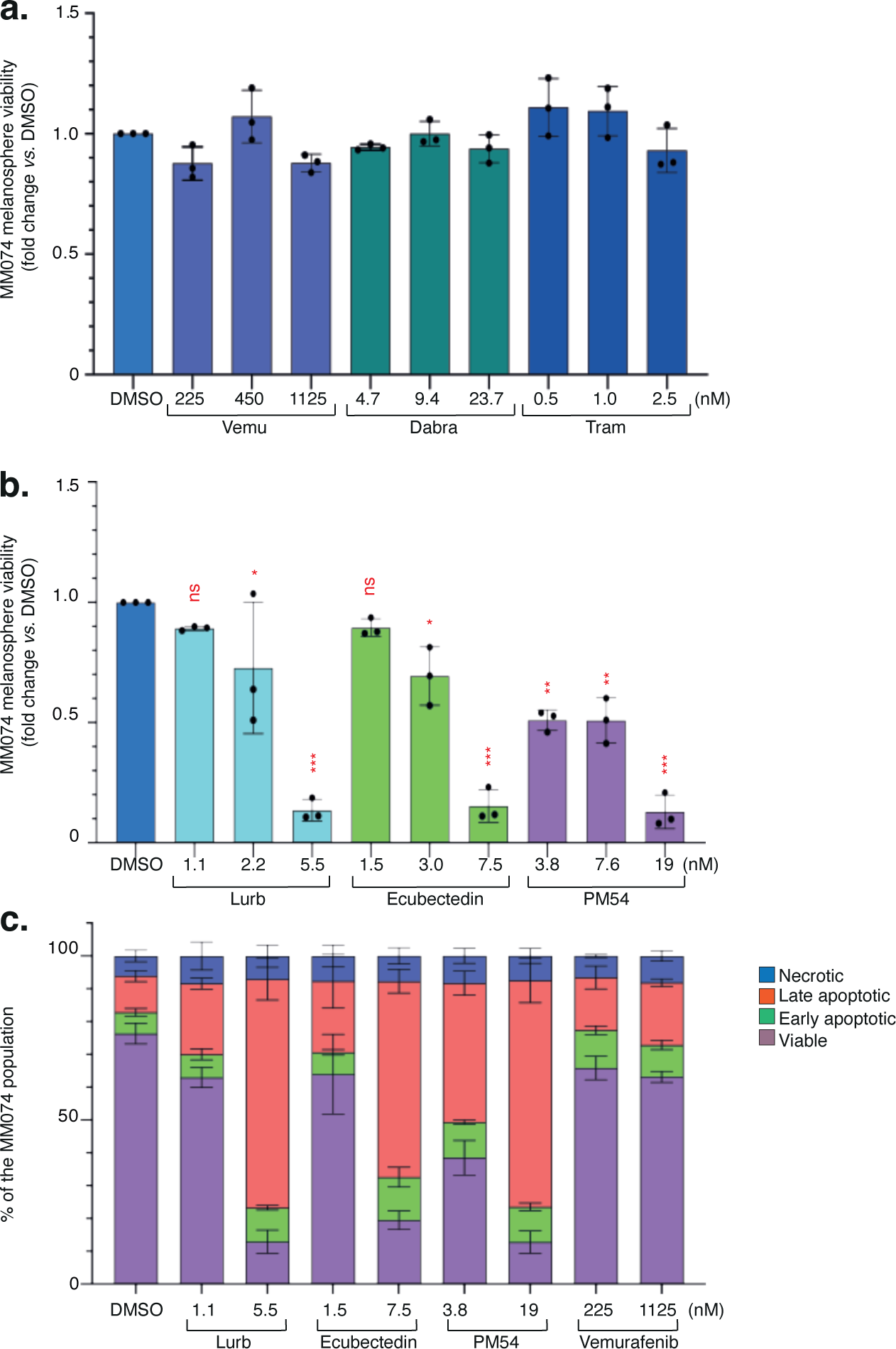
Melanospheres have high sensitivity to synthetic ecteinascidins. **a-b.** MM074 melanospheres were treated with drugs as indicated for 72 hours, and cell viability was measured with CellTiter-Glo assay. Results are shown as mean values of viability *vs.* DMSO +/− SD for three biological triplicates. Ordinary one-way ANOVA using Dunnett’s multiple comparisons test was used to determine the p-values. **c.** MM074 melanospheres were treated with either vehicle (DMSO), lurbinectedin, ecubectedin, PM54 or vemurafenib as indicated for 72 hours. Apoptosis was studied by flow cytometry with Annexin V-APC and propidium iodide staining. Results are shown as mean values +/− SD for three biological triplicates.

**Supplemental Figure 4:**
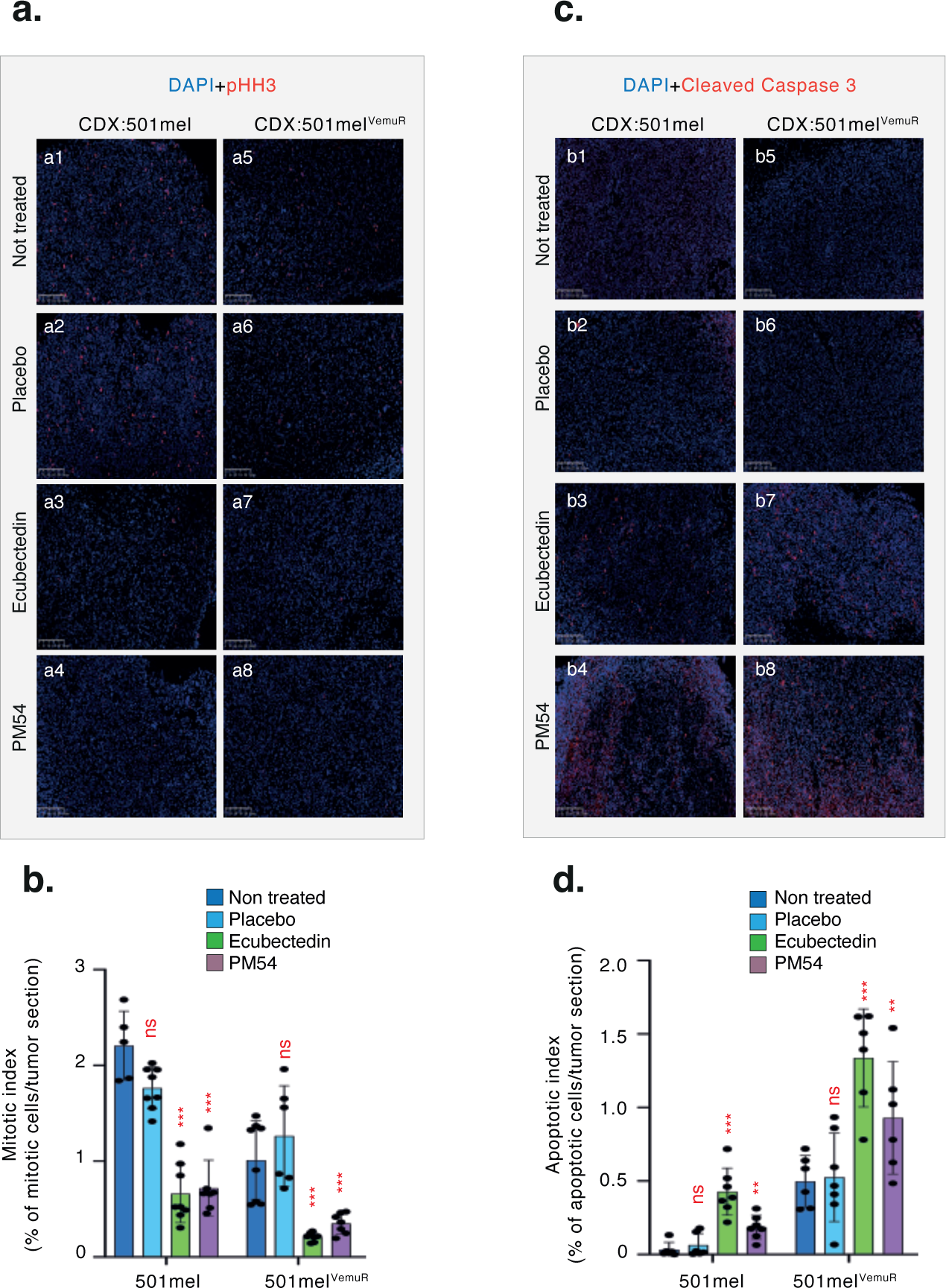
Synthetic ecteinascidins inhibit cell proliferation and induce apoptosis *in vivo*. **a-c.** Representative images of pHH3-positive cells **(a)** and cleaved caspase-3-positive cells **(c)** in tumor sections of 501mel or 501mel^VemuR^ CDX models after a 24-hour treatment with a one-time dose of Placebo, ecubectedin or PM54 at 1.2 mg/kg. **b.** Quantification of the mitotic index (% of pHH3-positive cells/tumour section(n=3)) is shown as mean values +/− SD for tumour sections. Ordinary one-way ANOVA using Dunnett’s multiple comparisons test was used to determine the p-values (*vs.* non treated). **c**. Quantification of the apoptotic index (% of cleaved caspase-3-positive cells/tumor section(n=3)) is shown as mean values +/− SD for tumour sections. Ordinary one-way ANOVA using Dunnett’s multiple comparisons test was used to determine the p-values (*vs.* non treated).

**Supplemental Figure 5:**
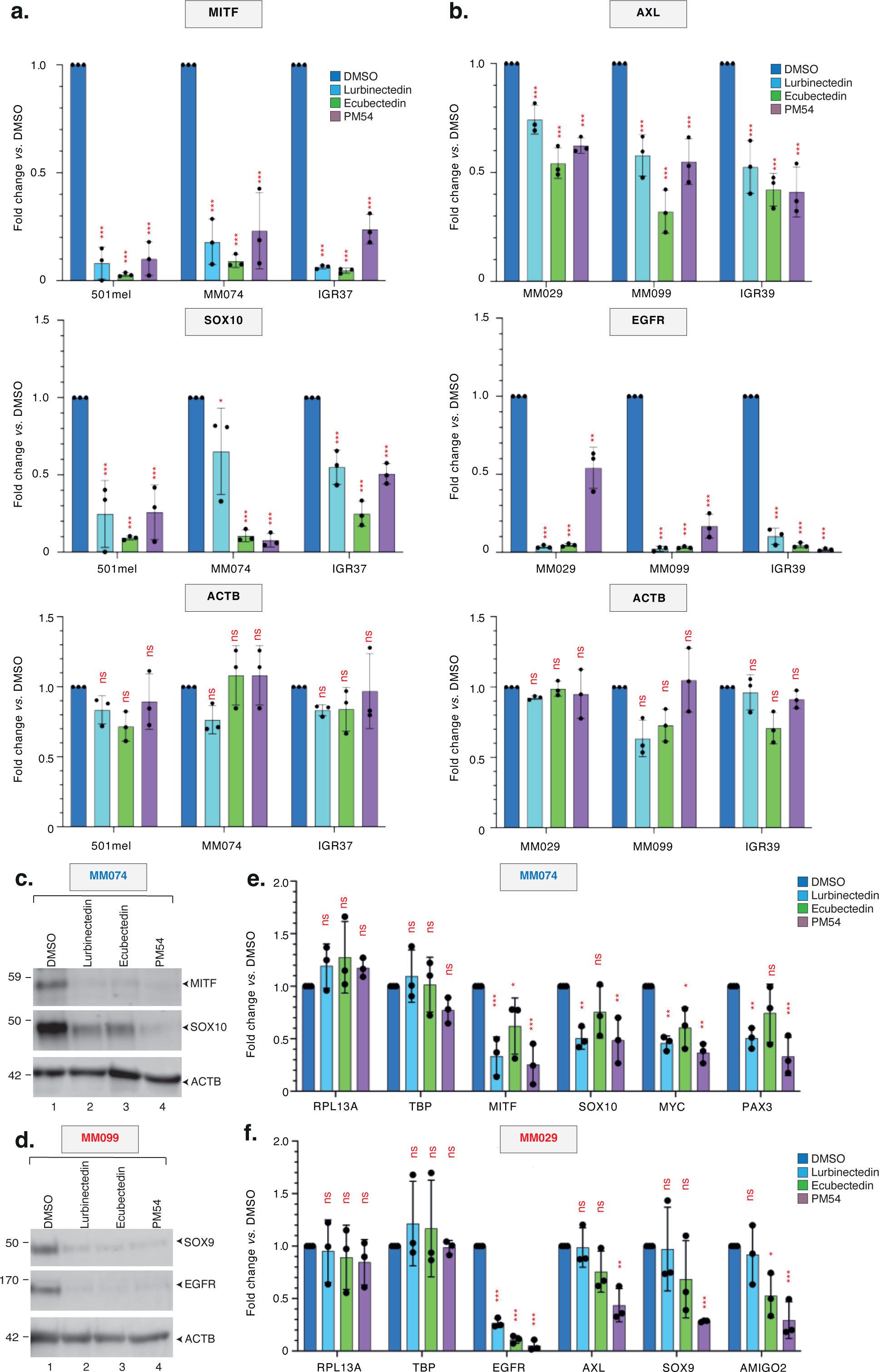
Synthetic ecteinascidins impair crucial cancer-promoting melanoma genes. **a.** qRT-PCR analysis showing average 18S-normalized expression of MITF, SOX10 and ACTB in the differentiated 501mel, MM074 and IGR37 cells treated with either vehicle (DMSO), lurbinectedin, ecubectedin or PM54 (5xIC50 concentration, 12 hours). Error bars indicate mean values + SD for three biological triplicates. Ordinary one-way ANOVA using Dunnett’s multiple comparisons test was used to determine the p-values (*vs.* DMSO). **b.** qRT-PCR analysis showing average 18S-normalized expression of AXL, EGFR and ACTB in the undifferentiated MM029, MM099 and IGR39 treated with either vehicle (DMSO), lurbinectedin, ecubectedin or PM54 (5xIC50 concentration, 12 hours). Error bars indicate the mean values +/− SD for three biological triplicates. Ordinary one-way ANOVA using Dunnett’s multiple comparisons test was used to determine the p-values (*vs.* DMSO). **c-d.** The differentiated MM074 **(c)** or undifferentiated MM099 **(d)** cells were treated with either vehicle (DMSO), lurbinectedin, ecubectedin or PM54 (5xIC50 concentration, 24 hours). Protein lysates were immuno-blotted for proteins as indicated. Molecular mass of the proteins is indicated (kDa). **e-f.** qRT-PCR analysis showing average 18S-normalized gene expression in the differentiated MM074 **(e)** and undifferentiated MM029 **(f)** melanospheres treated with either vehicle (DMSO), lurbinectedin, ecubectedin or PM54 (5xIC50 concentration, 24 hours). Error bars indicate mean values + SD for three biological triplicates. Ordinary one-way ANOVA using Dunnett’s multiple comparisons test was used to determine the p-values (*vs.* DMSO).

**Supplemental Figure 6:**
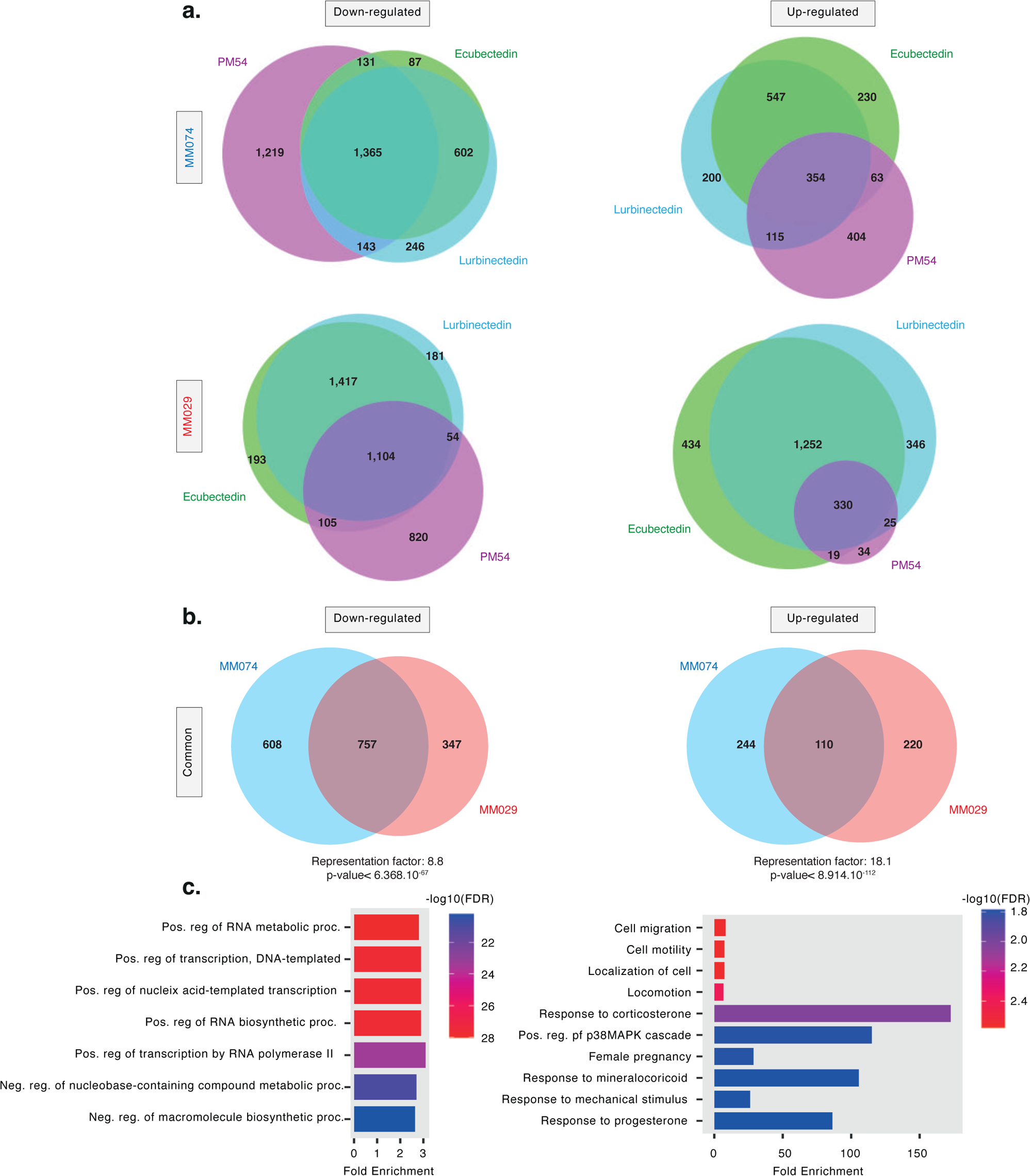
Comparison of down-and up-regulated genes in melanoma cell subtypes following treatment with synthetic ecteinascidins. **a.** Venn diagram between significantly down-regulated (left) and up-regulated (right) genes identified by RNA-seq in differentiated MM074 (top) and undifferentiated MM029 (bottom) upon treatment with either lurbinectedin, ecubectedin or PM54 (10xIC50 concentration, 8 hours). **b.** Venn diagram between genes identified by RNA-seq as being commonly down-regulated (left) or up-regulated (right) in both differentiated MM074 and undifferentiated MM029 by the three synthetic ecteinascidins. Representation factor and hypergeometric p-values are represented. **c.** Gene ontology (Biological process) analysis of the 757 genes significantly down-regulated (left) and 110 genes significantly up-regulated in both MM074 and MM029 by the three synthetic ecteinascidins, as identified in **(b)**. The histogram shows the top deregulated biological pathways according to the FDR and fold enrichment.

**Supplemental Figure 7:**
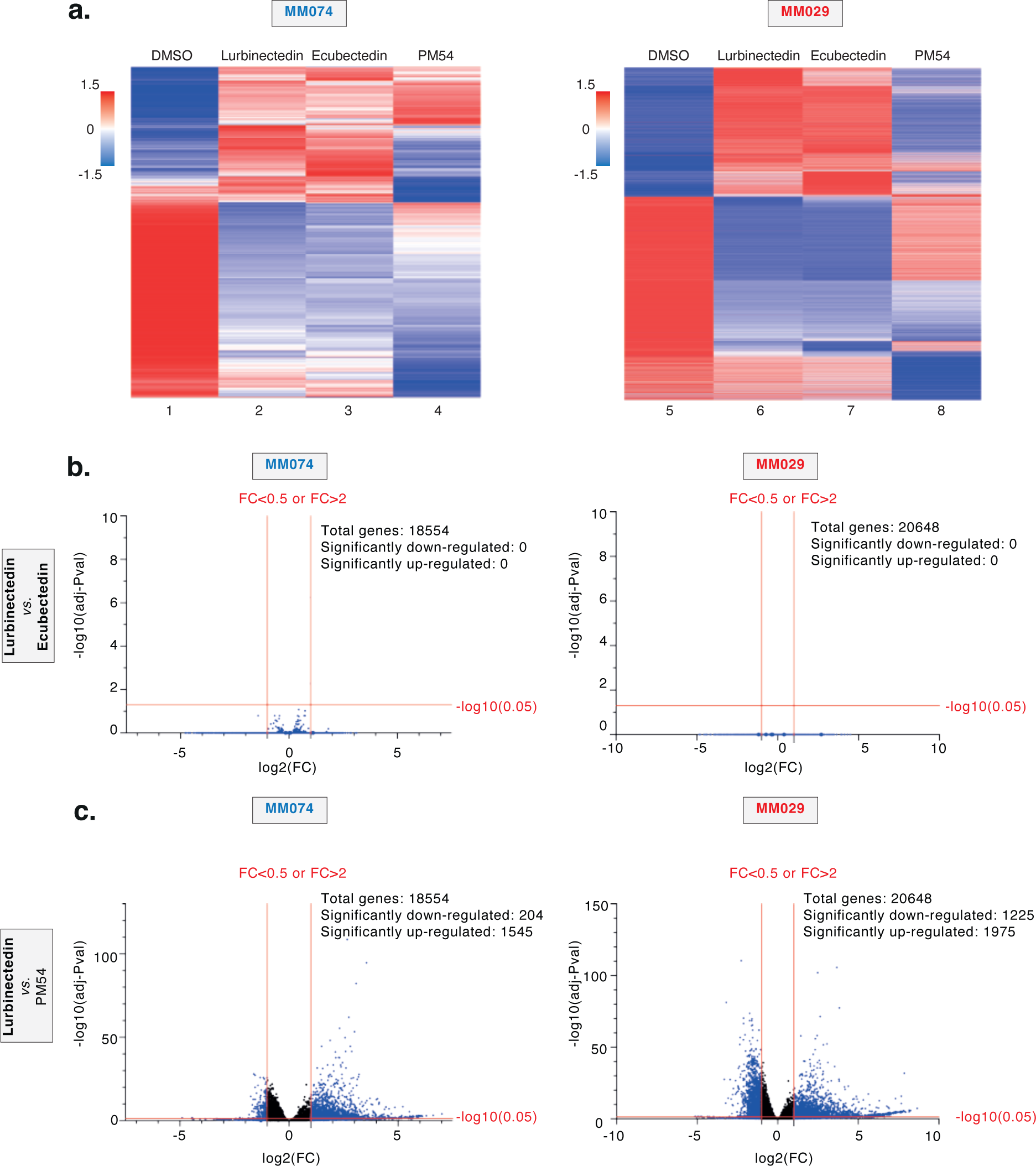
PM54 differentially affects gene expression compared to lurbinectedin and ecubectedin. **a.** Heatmap depicting all deregulated genes from either lurbinectedin, ecubectedin or PM54 treatments (10xIC50 concentration, 8 hours), in MM074 cells (left) or MM029 cells (right). RPKM values are represented as z-score. **b-c.** Volcano plots showing differentially expressed genes between lurbinectedin and either ecubectedin **(b)** or PM54 **(c)** treatment in MM074 (left) and MM029 (right) as determined by RNA-seq. Deregulated genes were defined as genes with log2(Fold change) > 1 or < −1 and adjusted P-value < 0.05.

**Supplemental Figure 8:**
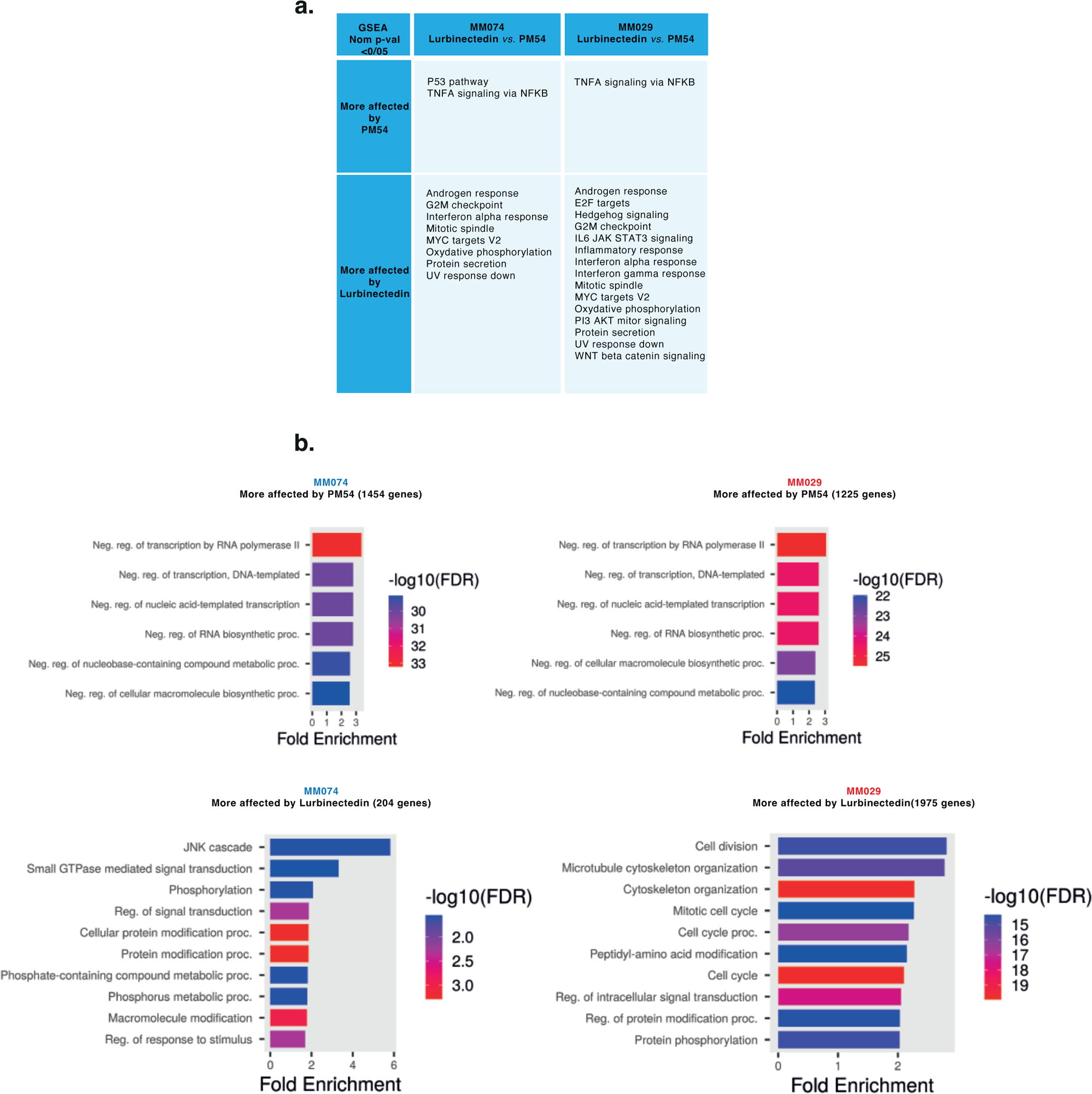
PM54 more directly affects genes encoding for transcription regulators. **a.** GSEA analysis of differentially deregulated genes in MM074 and MM029 cells treated either with lurbinectedin or PM54, determined by RNA-seq as described in Figure 5. **b.** GO (Biological process) analysis of the genes significantly up-regulated (up) or down-regulated (bottom) in MM074 (left) and MM029 (right) cells treated with lurbinectedin *vs.* PM54. The histogram shows the top deregulated biological pathways according to the FDR and fold enrichment.

**Supplemental Figure 9:**
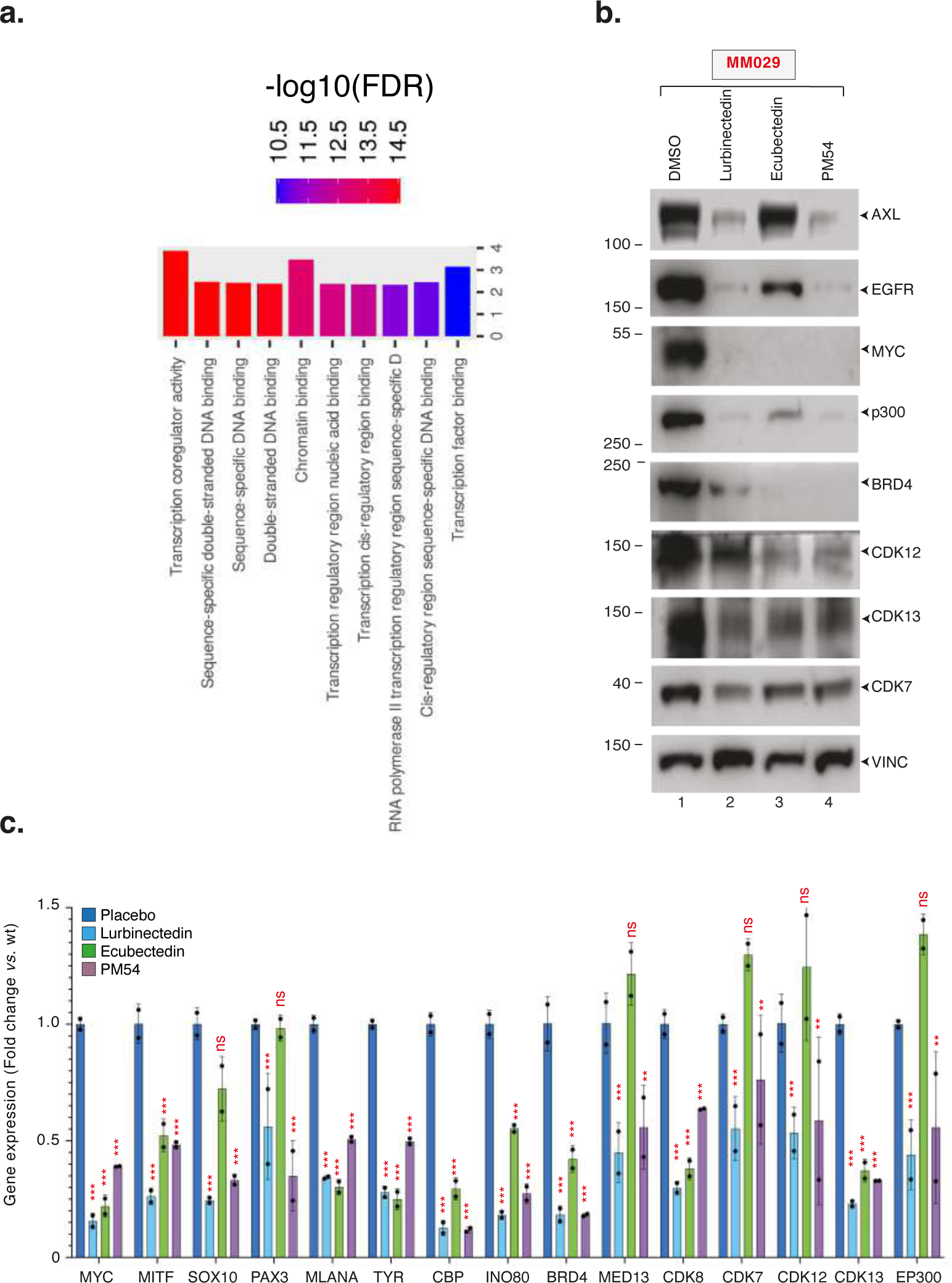
Synthetic ecteinascidins inhibit expression of genes coding for transcription factors/coactivators. **a.** GO (Molecular Function) analysis of the common set of 642 genes that underwent significant down-regulation upon lurbinectedin exposure of SKCM, SCLC and Non-SCLC cells. The histogram shows the top deregulated biological pathways according to the FDR and fold enrichment. **b.** Undifferentiated MM029 melanoma cells were treated with synthetic ecteinascidins as indicated (5xIC50, 24 hours) and protein lysates were immuno-blotted for proteins as indicated. Molecular mass of the proteins is indicated (kDa). **c.** CDXs from 501mel (n=3) were treated with a single dose of lurbinectedin, ecubectedin or PM54 at 1.2 mg/kg and tumors were collected 12 hours later. qRT-PCR analysis shows average placebo-normalized expression of the indicated genes (+/−SD). Ordinary one-way ANOVA using Dunnett’s multiple comparisons test was used to determine the p-values (*vs.* Placebo).

**Supplemental Figure 10:**
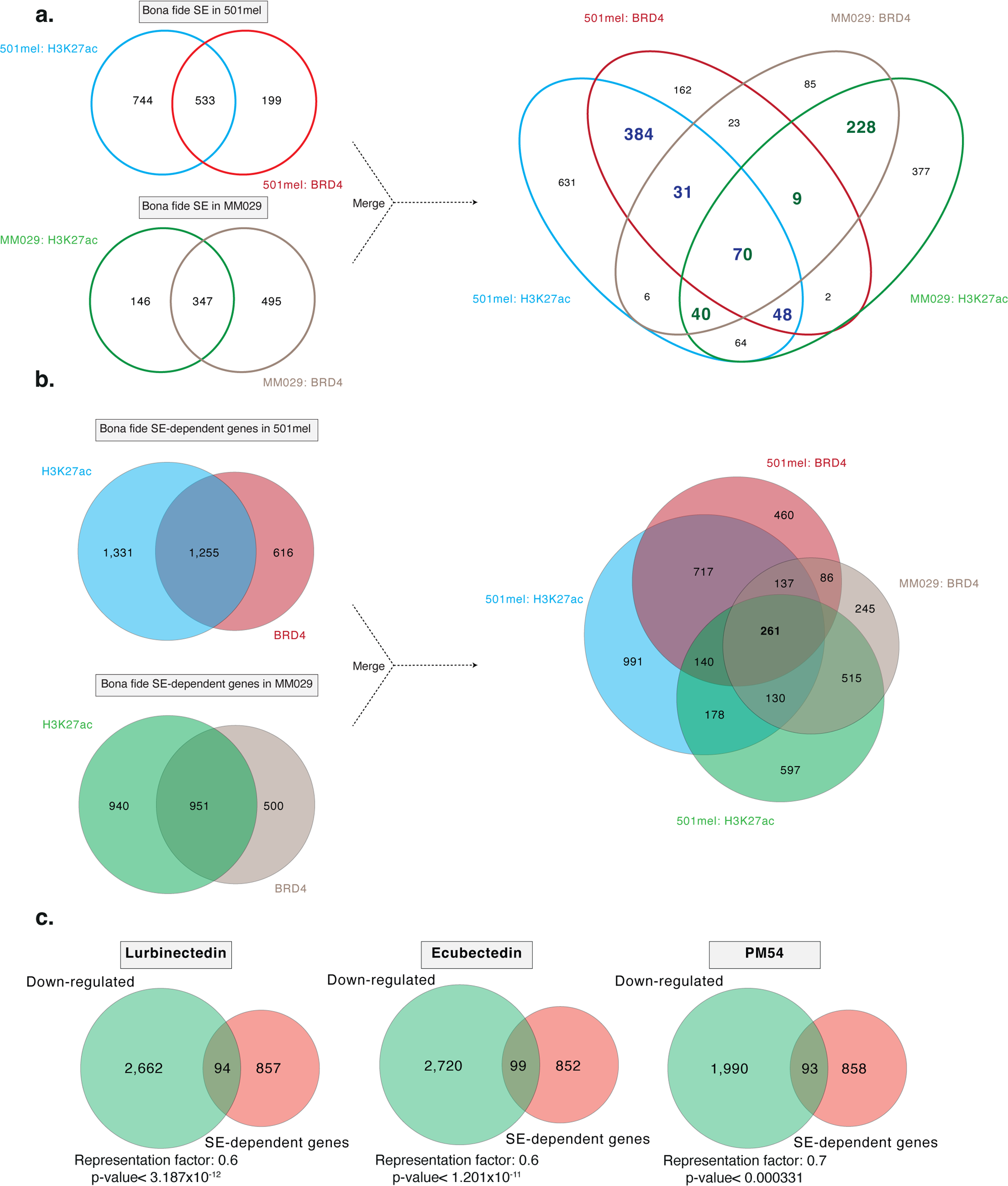
Identification of SEs in differentiated and undifferentiated melanoma cells. **a. Left panel**; Venn diagram between SEs identified by ROSE using either the H3K27ac- or BRD4-occupied sites in differentiated 501mel or undifferentiated MM029 cells. We defined as ‘bona fide’ SEs bound by both H3K27ac and BRD4. **Right panel**; the two Venn diagrams were merged. **b. Left panel**; Venn diagram between SE-dependent genes identified by ROSE using either the H3K27ac- or BRD4-occupied sites in differentiated 501mel or undifferentiated MM029 cells. We defined as ‘bona fide’ SE-dependent genes if their SEs are bound by both H3K27ac and BRD4. **Right panel**; the two Venn diagrams were merged. **c.** Venn diagram showing the overlap of genes downregulated by synthetic ecteinascidins, as indicated, in MM029 cells (10xIC50 concentration, 8 hours) and bona fide SE-dependent genes identified in MM029 cells. Representation factor and hypergeometric p-value are indicated.

**Supplemental Figure 11:**
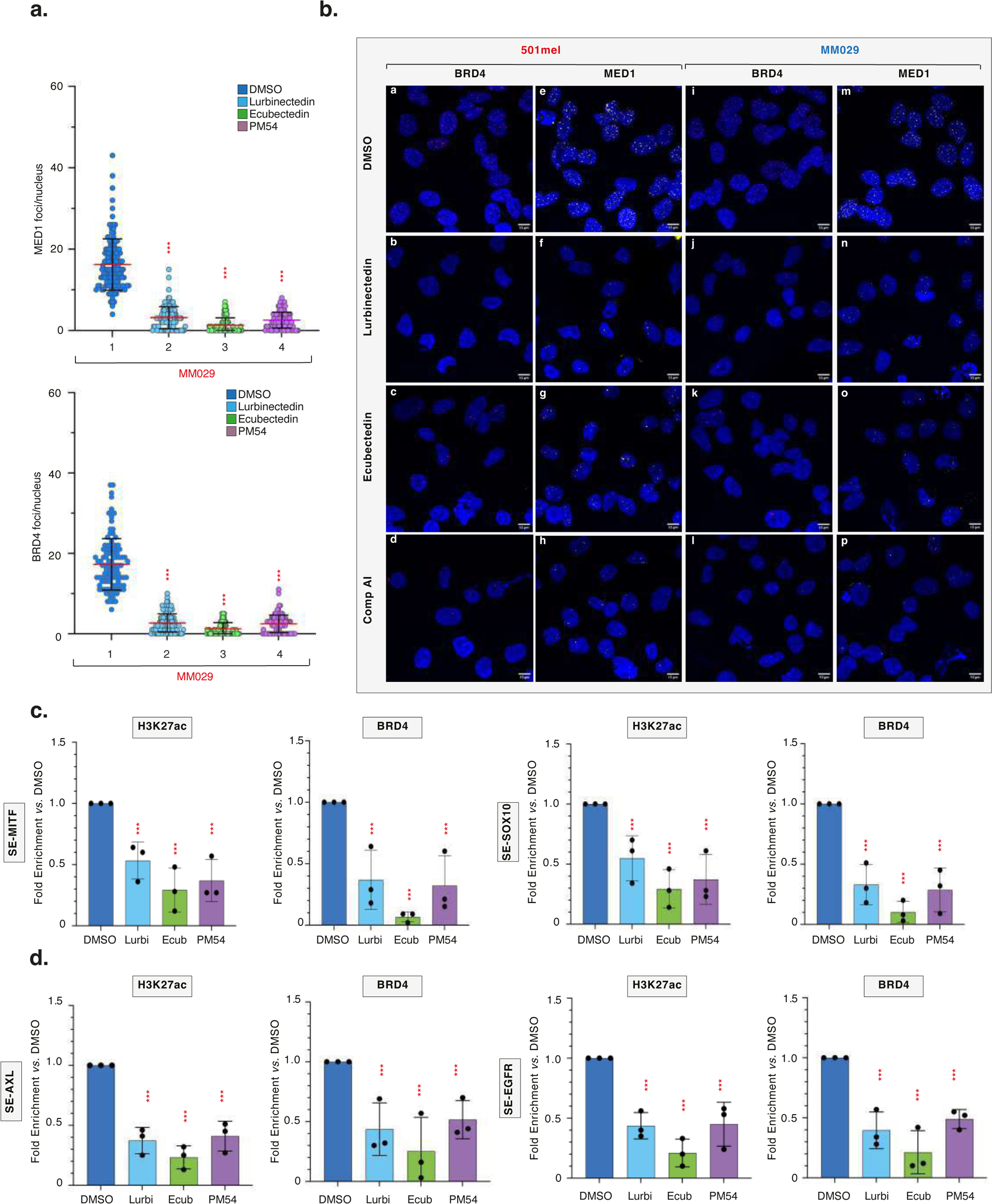
Synthetic ecteinascidins decommission SEs in melanoma. **a.** The numbers of MED1 (top) and BRD4 (bottom) foci per nucleus observed in MM029 cells following treatment with DMSO or synthetic ecteinascidins are shown +/− SD. Red bars indicate mean integrated density. One-way ANOVA with post-hoc Tukey adjustment comparisons were used to determine the p-values (*vs.* DMSO). **b.** Representative confocal images of 501mel or MM029 melanoma cells mock- or synthetic ecteinascidin-treated (5xIC50, 24 hours). Cells were immunostained with anti-BRD4 (red) or anti-MED1 (white) antibodies. Images of the cells were obtained with the same microscopy system and constant acquisition parameters for a given staining. **c.** ChIP/qRT-PCR monitoring the fold enrichment of H3K27ac mark (left) or BRD4 protein (right) at the SEs regulating MITF (left) or SOX10 (left) (+/− SD) in differentiated 501mel cells mock- or synthetic ecteinascidin-treated (5xIC50, 24 hours). One-way ANOVA with post-hoc Tukey adjustment comparisons were used to determine the p-values (*vs.* DMSO). **d.** ChIP/qRT-PCR monitoring the fold enrichment of H3K27ac mark or BRD4 protein at the SEs regulating AXL (left) and EGFR (right) (+/− SD) in undifferentiated MM029 cells mock- or synthetic ecteinascidin-treated (5xIC50, 24 hours). One-way ANOVA with post-hoc Tukey adjustment comparisons were used to determine the p-values (*vs.* DMSO).

**Supplemental Figure 12:**
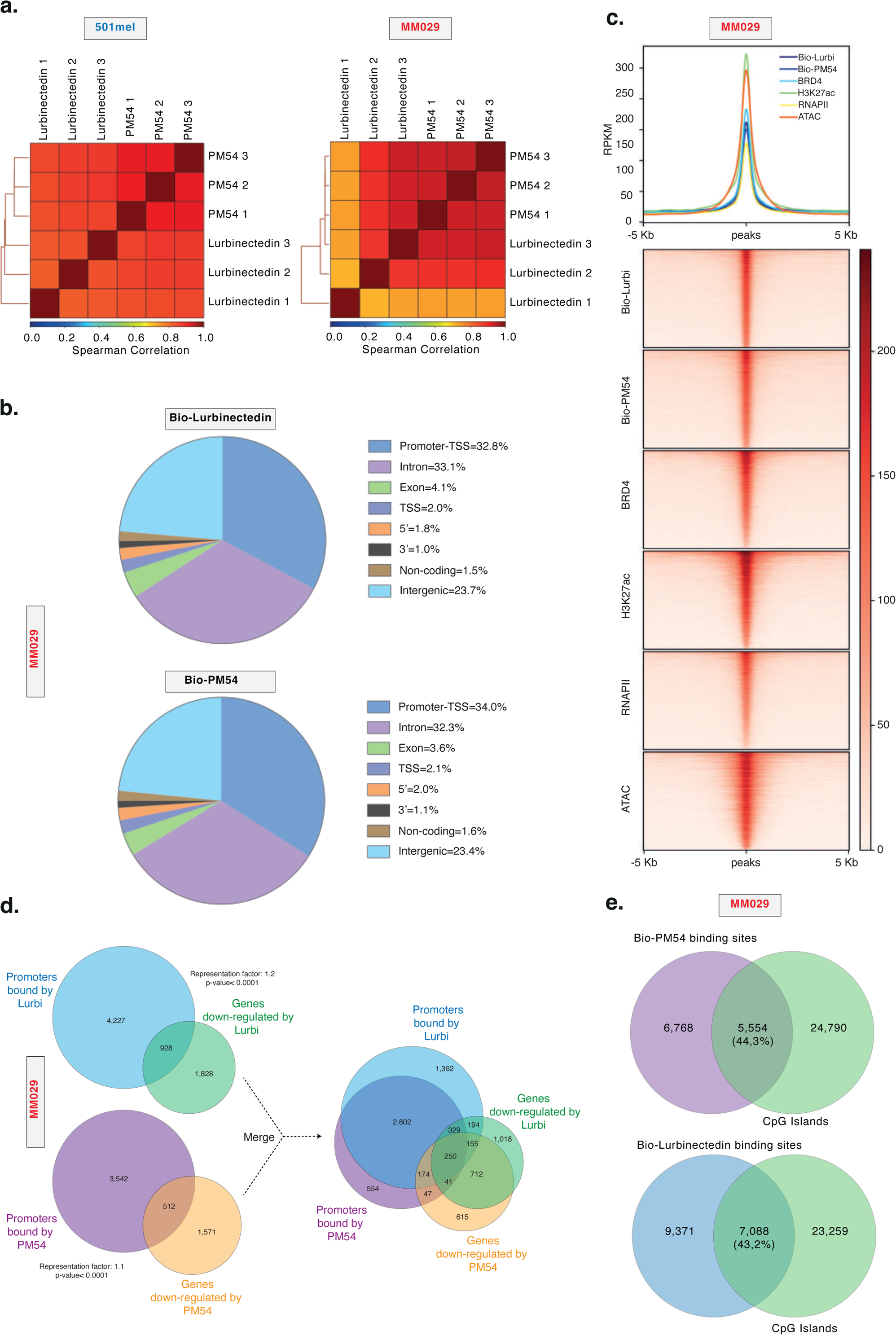
Synthetic ecteinascidins engage with transcriptionally active genomic regions in melanoma cells. **a.** Spearman correlation between triplicates of chemical-mapping analysis of Bio-lurbi and Bio-PM54 in 501mel (left) or MM029 (right) cells. **b.** Pie chart showing the distribution of Bio-lurbi-(top) and Bio-PM54-(bottom) annotated peaks (in percentage) all over the genome (hg19) in undifferentiated MM029 cells. **c. Upper panel;** Metaplot distribution of Bio-lurbi, Bio-PM54, BRD4, RNAPII, H3K27ac enrichment and ATAC-Seq signals in a +/−5kb window around the occupied DNA binding sites of Bio-lurbi in undifferentiated MM029 cells. **Lower panel**; Heatmap profiles representing the read density clusterings obtained with seqMINER for the DNA-occupied sites of Bio-lurbi in undifferentiated MM029 cells relative to Bio-PM54, BRD4, RNAPII, H3K27ac enrichments and ATAC-Seq signals. Peak order is determined by Bio-lurbi and identical for all clusterings. **d. Left panel;** Venn diagram between promoters bound by Bio-lurbi or Bio-PM54 and genes down-regulated by Lurbinectedin or PM54 in MM029 cells. **Right panel**; the two Venn diagrams were merged. Representation factor and hypergeometric p-value are indicated. **e.** Venn diagram between Bio-lurbi (top) and Bio-PM54 (bottom) binding sites identified in MM029 cells and human CpG Islands.

**Supplemental Figure 13:**
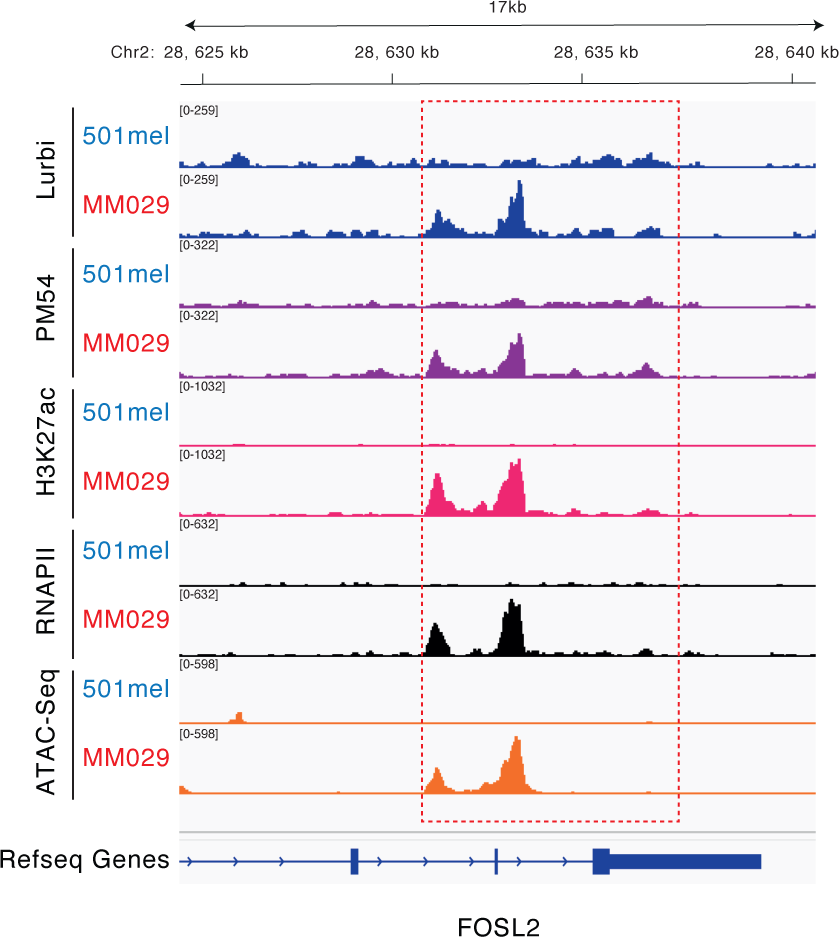
Synthetic ecteinascidins bind to the SE regulating FOSL2 expression. Gene tracks of Bio-lurbi, Bio-PM54, RNAPII, H3K27ac occupancy and ATAC-seq signals at the *FOSL2* locus in 501mel or MM029 cells. This gene is only expressed and bound Bio-lurbi and Bio-PM54 in undifferentiated MM029 melanoma cells. The red square indicates a portion of the *FOSL2* SE.

**Supplemental Figure 14:**
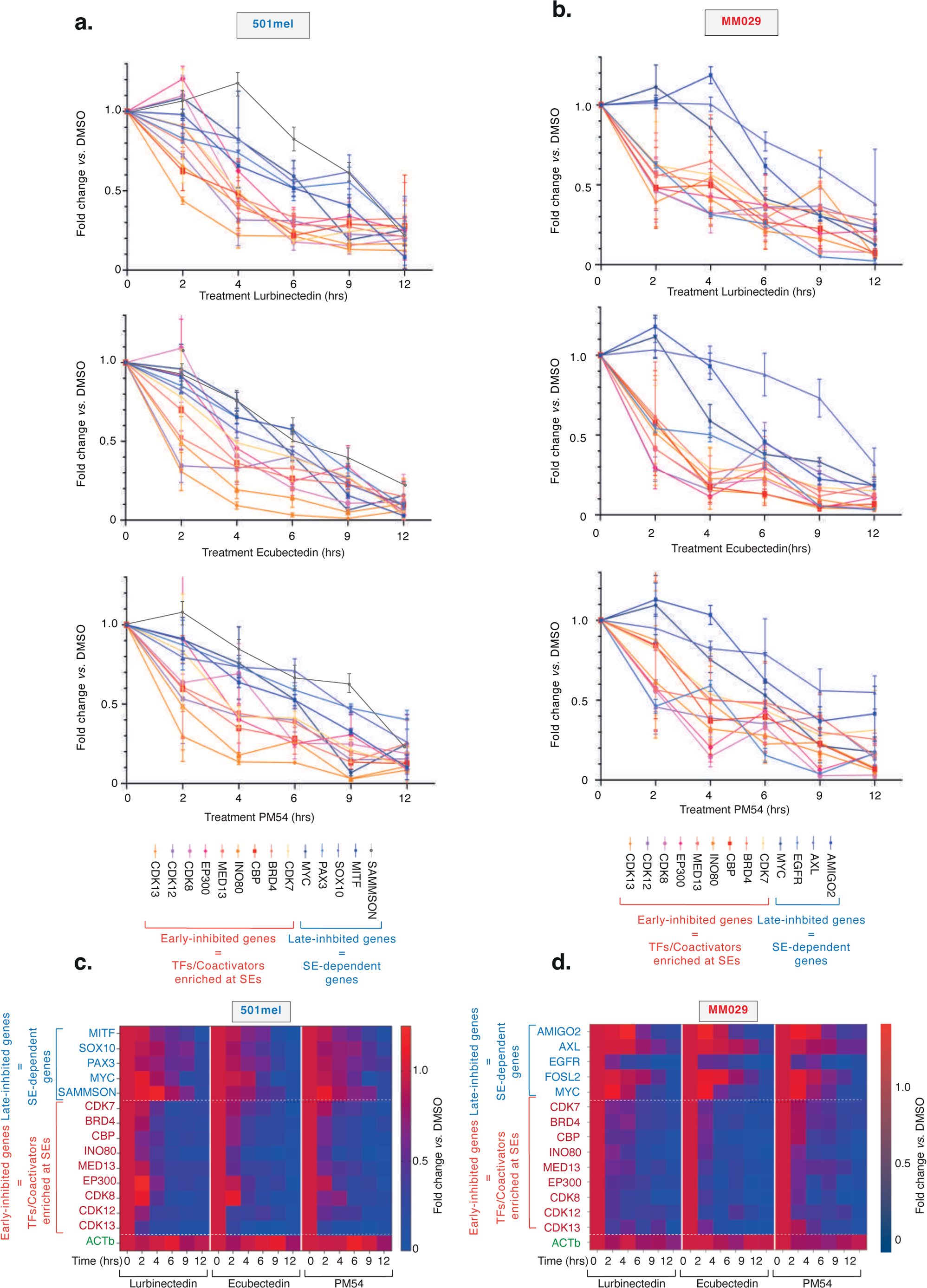
Synthetic ecteinascidins affect transcription factors/coactivators and specific SE-dependent genes in SKCM. **a-b.** qRT-PCR analysis showing average 18S-normalized expression of the indicated genes in the differentiated 501mel **(a)** or undifferentiated MM029 melanoma cells **(b)** following treatment with Lurbinectedin, Ecubectedin or PM54 (5xIC50 concentration) for the indicated period of time. Error bars indicate the mean values +/− SD for three biological triplicates. Results are shown as relative expression compared to mock-treated cells. **c-d**. Heatmap showing average 18S-normalized expression of the indicated genes in 501mel **(d)** or MM029 **(d)** cells treated with either Lurbinectedin, Ecubectedin or PM54 (5xIC50 concentration) for the indicated period of time. Results were obtained by RT-qPCR performed in (a) and are shown as relative expression compared to DMSO-treated cells. ACTb is a housekeeping gene.

**Supplemental Figure 15:**
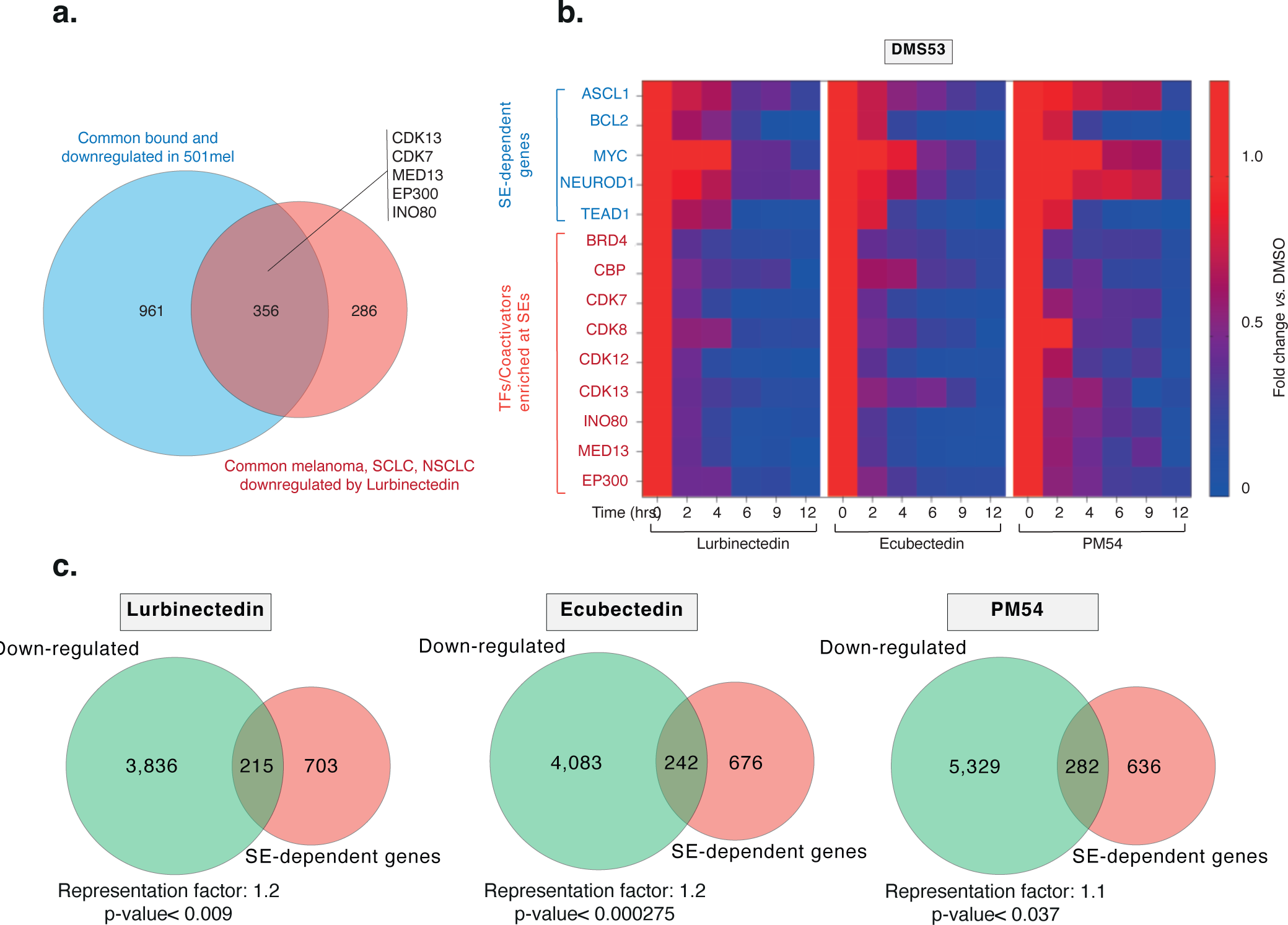
Synthetic ecteinascidins affect transcription factors/coactivators and specific SE-dependent genes in SCLC. **a.** Venn diagram showing the overlap of genes bound and downregulated by ecteinascidins in 501mel cells, and genes commonly downregulated in SKCM, SCLC and NSCLC. **b.** Venn diagram showing the overlap of genes downregulated by ecteinascidins, as indicated, in DMS53 SCLC cells (10xIC50 concentration, 8 hours) and SE-dependent genes identified in these cells^77^. Representation factor and hypergeometric p-value are indicated. **c.** Heatmap showing average 18S-normalized expression of the indicating genes in SCLC DMS53 cells treated with ecteinascidins (5xIC50 concentration) for the indicated period of time. Results were obtained by RT-qPCR and are shown as relative expression compared to DMSO-treated cells.

